# The topology of DNA entrapment by cohesin rings

**DOI:** 10.1101/495762

**Authors:** Christophe Chapard, Robert Jones, Till van Oepen, Johanna C Scheinost, Kim Nasmyth

## Abstract

Cohesin entraps sister DNAs within tripartite rings created by pairwise interactions between Smc1,Smc3, and Scc1. Because the ATPase heads of Smc1 and Smc3 can interact with each other, cohesin rings in fact have the potential to form a variety of sub-compartments. Using in vivo cysteine crosslinking,we show that when Smc1 and Smc3 ATPases are engaged in the presence of ATP (E heads)cohesin rings generate a “SMC (S) compartment” between hinge and E heads and a “kleisin (K)compartment” between E heads and their associated kleisin subunit. Upon ATP hydrolysis, cohesin’s heads associate with each other in a very different mode, in which their signature motifs and their coiled coils are closely juxtaposed (J heads), creating alternative S and K compartments. We show that all four sub-compartments exist in vivo, that acetylation of Smc3 during S phase is accompanied by an increase in the ratio of J to E heads, and that sister DNAs are entrapped in J-K but not E-K compartments or in either type of S compartment.

## Introduction

The cohesin complex not only holds sister chromatids together in post-replicative proliferating cells (Guacci et al., 1997; Michaelis et al., 1997) but also organizes the topology of chromatin fibres during interphase (Rao et al., 2017). The former involves interactions between different DNA molecules that must be stable for very extended periods of time, possibly years in the case of meiotic cells (Hunt and Hassold, 2010), while the latter involves transient long range interactions between sequences from the same DNA molecule that organize chromosomal DNAs into chromatid-like threads with loops emanating from a central axis (Klein et al., 1999; Tedeschi et al., 2013). Given these differences, the actual mechanisms are likely to be different. It has been suggested that sister chromatid cohesion is mediated by co-entrapment of sister DNAs within a tripartite ring formed by pairwise interactions between cohesin’s Smc1, Smc3, and kleisin (Scc1) subunits (Haering et al., 2002) while chromatid-like structures during interphase are created by a DNA translocase associated with cohesin that progressively extrudes ever longer loops of DNA (Nasmyth, 2001), an activity thought to be responsible for creating the Topologically Associated Domains (TADs) observed using HiC (Fudenberg et al., 2016; Haarhuis et al., 2017; Rao et al., 2017; Sanborn et al., 2015; Schwarzer et al., 2017; Wutz et al., 2017). Whether loop extrusion also involves entrapment of DNAs within cohesin rings is not known.

Understanding the detailed topology of cohesin’s interactions with DNA while it confers cohesion or undergoes loop extrusion is therefore crucial to understanding these two rather different functions. A key aspect of this topology is the potential for DNAs to be entrapped inside a variety of compartments within rings created by multiple interactions between cohesin’s Smc1, Smc3, and Scc1 subunits. Smc1 and Smc3 are rod shaped proteins with dimerization domains at one end and ABC-like ATPase domains at the other, connected by 50 nm long coiled coils. Dimerization creates V-shaped Smc1/Smc3 heterodimers with a hinge at their junction and ATPases at their vertices (Haering et al., 2002; Haering et al., 2004). The association of Scc1’s N-and C-terminal domains with the coiled coil emerging from Smc3’s ATPase (its neck) and the base of Smc1’s ATPase respectively creates a huge SMC-kleisin (SK) ring (Gligoris et al., 2014; Haering et al., 2004). Additional interactions between Smc1 and Smc3 in the vicinity of their ATPase heads may divide the large ring created by joining Smc hinge and Smc/kleisin interfaces into two sub-compartments as described in the present study, namely a “SMC (S) compartment” created by the Smc1/3 hinge and Smc1/3 head interactions and a “kleisin (K) compartment” defined by Smc1/3 head interactions and interactions of each ATPase head with the N- and C-terminal domains of Scc1.

Work on a related Smc/kleisin complex from B. subtilis suggests that Smc heads in fact interact in two very different ways (Diebold-Durand et al., 2017; Minnen et al., 2016). The first involves the interaction between ATPs bound to one head (Smc1) with signature motifs on its partner (Smc3) and vice versa, creating a complex that sandwiches a pair of ATP molecules (Arumugam et al., 2003; Lammens et al., 2004; Marcos-Alcalde et al., 2017). This ATP-induced Head Engagement (E) is a prerequisite to the hydrolysis of both ATP molecules that triggers disengagement and formation of a new state created by the rotation of both ATPase heads in a manner that juxtaposes their two signature motifs and their necks (signature motif Juxtapositon - J) (Diebold-Durand et al., 2017). This process may be facilitated by interactions between their coiled coils in the vicinity of a pronounced disruption known as the joint (Diebold-Durand et al., 2017). If the cohesin ring also undergoes a similar switch, which was one of the goals of the current study, then it could create five different types of compartments: S and K compartments associated with E and J heads as well as open SK rings in which neither heads nor coiled coils are juxtaposed (Fig. 1A).

**Figure 1.**
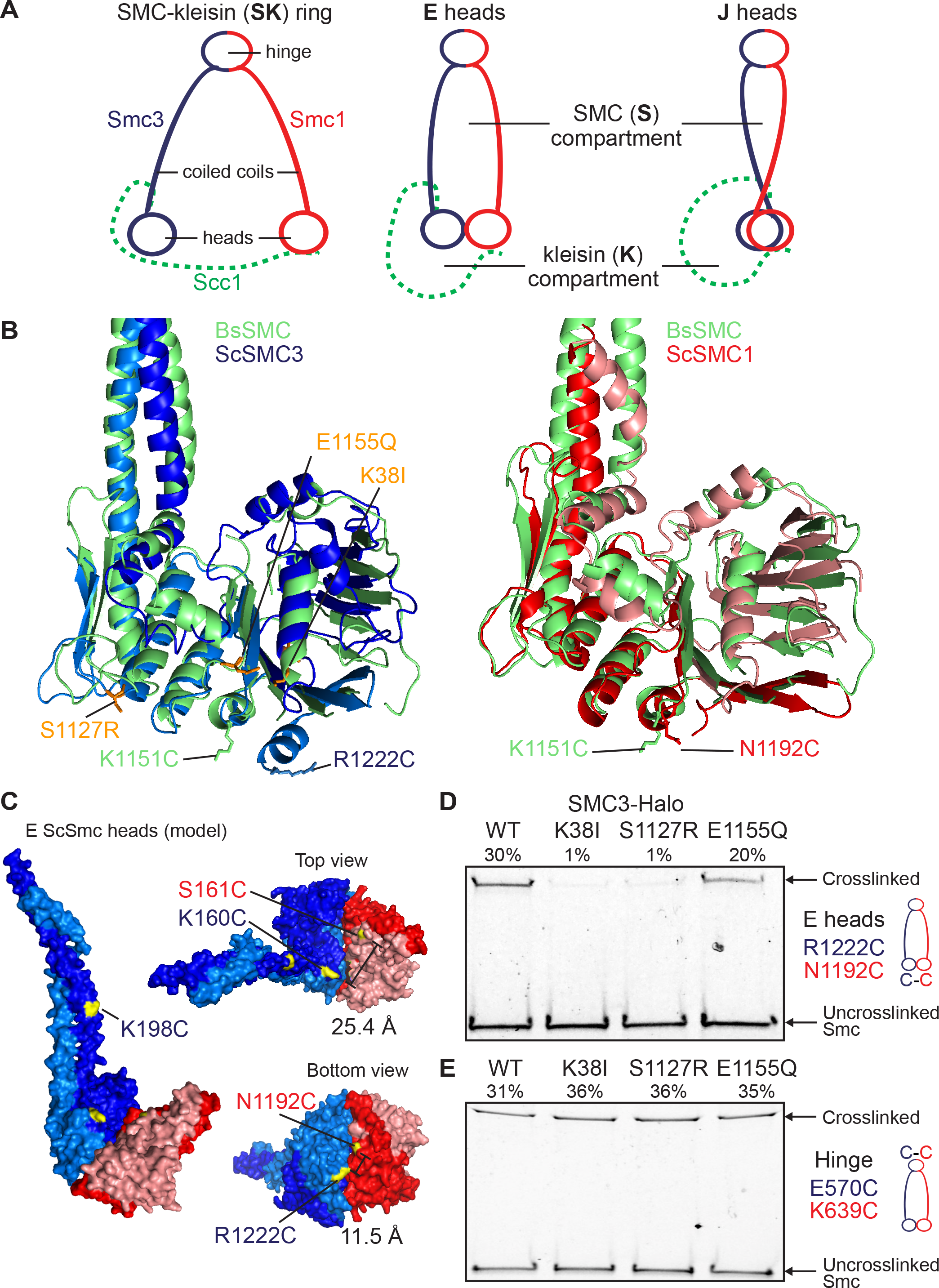
ATP dependent head engagement (E) state of Smc head domains. (A) Schematic representation of the cohesin compartments. (B) Structure alignment of Sc Smc3 head (PDB: 4UX3, blue) and Sc Smc1 head (PDB: 1W1W, red) to Bs Smc head (PDB: 3ZGX, green). Selected residues displaying efficient cross-linking when mutated to cysteine are marked (for B. subtilis, see Diebold-Durand et al., 2017). Residues associated with the Smc3 ATP binding mutant (K38I), the signature motif mutant (S1127R) and the ATP hydrolysis mutant (E1155Q) are displayed in orange. (C) Model of ATP-engaged Smc3/Smc1 heads in surface representation in front (left) and top and bottom views (right). ATP-engaged head dimer is constructed by superimposition of Smc1 head to one of Smc3 head homodimer (Gligoris et al, 2014). Distance between selected residues is given. (D-E) In vivo cysteine cross-linking of Smc1(Cys) proteins with Halo-tagged wild-type and ATPase mutant Smc3. Cross-linking of Smc1(N1192C) and Smc3(R1222C) E head residues (C) or Smc1(K639C) and Smc3(E570C) hinge residues (D) was performed in vivo using BMOE. Cell extracts were labelled with HaloTag-TMR ligand. Smc-HaloTag species were separated by SDS-PAGE and quantified by in-gel fluorescence. Percentage of cross-link efficiency is indicated.

Specific Smc/kleisin interactions have hitherto been detected in vivo using a bi-functional thiol-specific cross-linking reagent, BMOE, to induce rapid cross-linking between cysteine residue pairs inserted within individual ring interfaces (Gligoris et al., 2014). The results of these experiments imply that about 25% of cohesin rings are cross-linked simultaneously at all three Smc1/3 hinge, Smc3/Scc1, and Smc1/Scc1 interfaces (Gligoris et al., 2014). Chemical closure in this manner can then be exploited to detect DNA entrapment. Thus, entrapment of individual circular DNAs by chemically circularized cohesin rings leads to a modest retardation in their migration during gel electrophoresis even when all proteins have been denatured by heating in the presence of SDS (Haering et al., 2008). Likewise, co-entrapment of monomeric sister DNAs within chemically circularized cohesin causes them to migrate as dimers instead of monomers. Because they are catenated exclusively by cohesin rings, these sister DNA pairs are known as catenated dimers (CDs) (Gligoris et al., 2014). Analysis of numerous mutants has revealed a perfect correlation between the incidence of CDs and whether cells had established sister chromatid cohesion (Srinivasan et al., 2018). Thus, co-entrapment of sister DNAs within individual cohesin rings provides a mechanistic explanation for cohesion and for how cleavage of Scc1 by separase triggers sister chromatid disjunction at anaphase (Uhlmann et al., 2000).

These studies have not hitherto taken into account the possibility that DNAs are entrapped within the ring’s sub-compartments, namely S or K compartments associated with E or J heads. Indeed, it has been proposed on numerous occasions that cohesion is in fact conferred by entrapment within E-S compartments and that the interconnection of E heads by kleisin merely reinforces this entrapment (Elbatsh et al., 2016; Huber et al., 2016; Li et al., 2017; Murayama et al., 2018; Murayama and Uhlmann, 2015; Stigler et al., 2016; Uhlmann, 2009, 2016). Support for this notion stems from the observation that abolition of Smc3 de-acetylation by inactivation of the *HOS1* de-acetylase delays sister chromatid disjunction during anaphase despite efficient Scc1 cleavage (Li et al., 2017). If DNAs were in fact entrapped within E-S compartments, then cleavage of their coiled coils by separase should suppress the delayed disjunction, which is precisely what was found. Entrapment within the S or K compartments of complexes whose heads are engaged is likewise consistent with the claim that cohesion can be established by viable Smc1D1164E mutations that are supposedly incapable of hydrolysing ATP (Camdere et al., 2015; Camdere et al., 2018; Elbatsh et al., 2016).

To observe more precisely the nature of DNA entrapment by cohesin rings, we have inserted a series of cysteine pairs into Smc1 and Smc3 capable of detecting BMOE-induced cross-linking associated with head engagement, head juxtaposition, or interactions between Smc1 and Smc3 coiled coils in the vicinity of their joint regions. We find that individual minichromosome DNAs are entrapped in K compartments but never in S compartments of either type while sister DNAs are trapped inside J-K compartments. Strikingly, acetylation that accompanies formation of cohesion during S phase is more frequently associated with head juxtaposition than head engagement, implying that cohesion throughout the genome is also mediated by cohesin complexes with J heads.

## Results

### Using thiol-specific cross-linking to measure ATP-dependent head engagement

In absence of a crystal structure of Smc1/3 head heterodimers, we identified Smc1 and Smc3 residues predicted to reside within the engaged interface by aligning the structures of homodimeric Smc1 and Smc3 heads from S. cerevisiae, crystalized in the presence of ATP (Figs 1B and 1C) (Gligoris et al., 2014; Haering et al., 2004). To measure E head conformation in vivo, we sought amino acid pairs that were poorly conserved and could be replaced by cysteines without affecting cell viability (Fig 1C, S1A and S1B). Smc1N1192C and Smc3R1222C are 11.5Å apart in our model (Fig. 1C) and were crosslinked in vivo in a manner dependent on both cysteines and BMOE (Fig 1D and S1C). As predicted for cross-linking specific for E, that between Smc1N1192C and Smc3R1222C was reduced by Smc3 mutations that compromise ATP binding to Smc3 heads (K38I) or their interaction with ATP bound to Smc1 heads (S1127R) but not by a mutation that merely prevents ATP hydrolysis (E1155Q) (Figs. 1B and 1D). In contrast, none of these mutations had any effect on cross-linking between cysteine pairs embedded within the Smc1/Smc3 hinge interface (Fig. 1E). Interestingly, Smc1N1192 and Smc3R1222 are located in the same position as K1151 in B. subtilis (Bs) Smc, whose replacement by cysteine was used to measure E heads in that organism (Fig. 1B)(Minnen et al., 2016). In contrast to K1151C, whose cross-linking was infrequent in wild type B. subtilis and greatly increased by an EQ hydrolysis mutation, cross-linking between Smc1N1192C and Smc3R1222C was readily detected in otherwise wild type S. cerevisiae (Sc) cells and largely unaffected by Smc3E1155Q.

### An alternative conformation of Smc1 and Smc3 heads: juxtaposition of their signature motifs

To determine whether Smc1 and Smc3 ATPase heads also adopt a J conformation, we aligned Smc1 and Smc3 head structures with associated sections of coiled coil to a model of J Bs Smc heads (Diebold-Durand et al., 2017) (Fig 2A). This pinpointed Smc1S161 and Smc3K160 as the residues most likely to correspond to Bs SmcS152, whose replacement by cysteine permits BMOE induced J cross-linking (Diebold-Durand et al., 2017). The same approach identified Smc3K198 as the residue most likely to correspond to Bs SmcD193, whose replacement by cysteine gives rise to BMOE induced cross-linking between the coiled coils emerging from J Bs Smc heads. Because an equivalent structural alignment was not possible in the case of Smc1, we instead used sequence homology between its coiled coil and that of other Smcs (Fig. S1A) to identify Smc1K201C as a potential partner for Smc3K198C, assuming that Smc1 and Smc3 coiled coils interact with each other in a manner similar to those from J Bs Smcs. Neither Smc1S161 nor Smc3K160 are highly conserved and their replacement by cysteines had little or no effect on spore viability, even when combined as Smc1S161C Smc3K160C double mutants. The substitution by cysteine of both Smc1K201 and Smc3198 was likewise tolerated (Fig. S1B).

**Figure 2.**
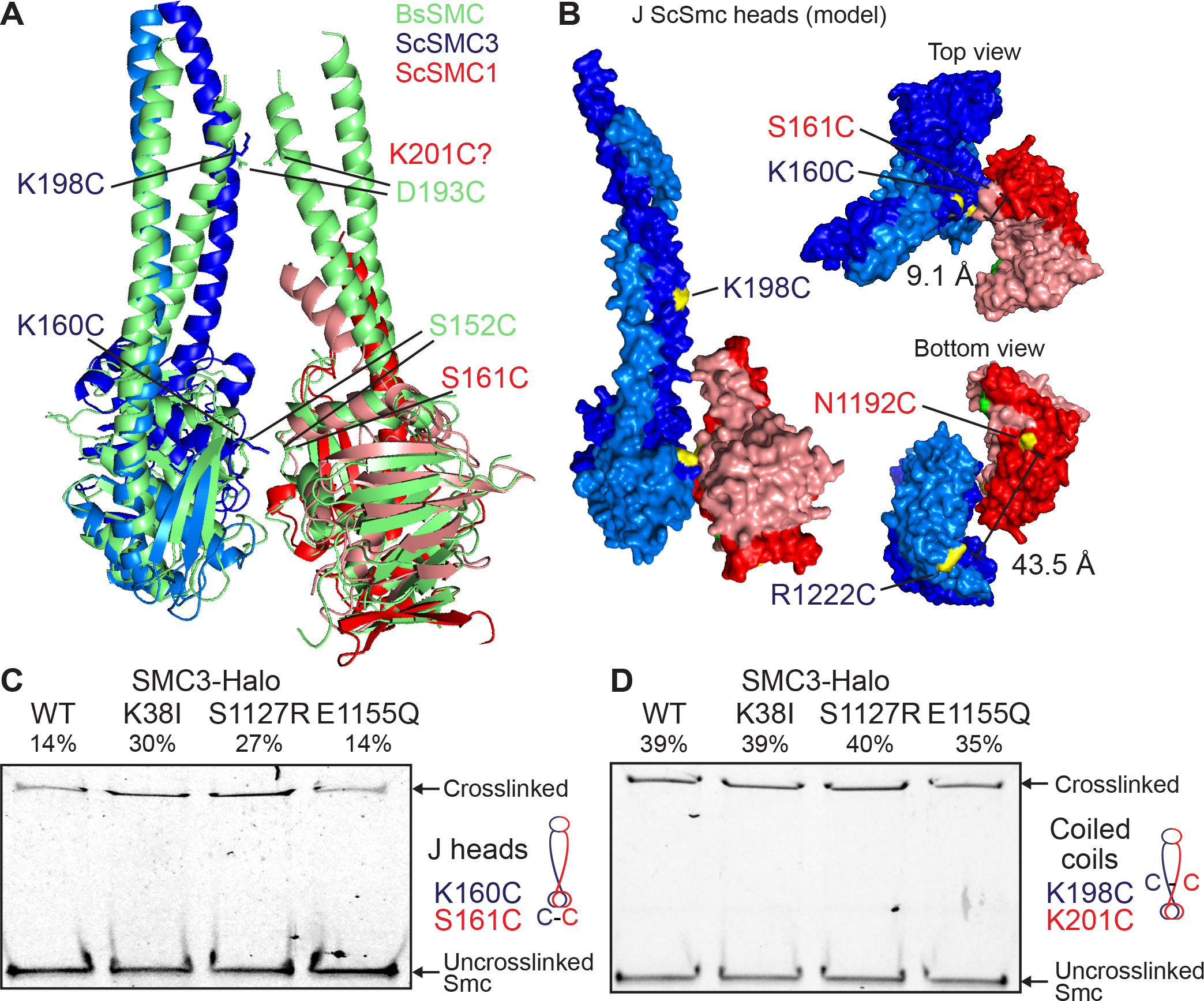
Signature motif juxtaposed (J) state of Smc head domains. (A) Structure alignment of Sc Smc3 head (PDB: 4UX3, blue) and Sc Smc1 head (PDB: 1W1W, red) to disengaged Bs Smc heads (PDB: 3ZGX, green). Selected residues displaying efficient cross-linking when mutated to cysteine are marked (for B. subtilis, see Diebold-Durand et al., 2017). (B) Model of J Smc3/Smc1 heads in surface representation in front (left) and top and bottom views (right). J head dimer is constructed by superimposition of Smc1/3 heads onto the rod aligned Bs Smc head model as in (A) (Diebold-Durand et al., 2017). Distance between selected residues is given. (C-D) In vivo cysteine cross-linking of Smc1(Cys) proteins with Halo-tagged wild-type and mutant ATPase Smc3. Cross-linking of Smc1(S161C) and Smc3(K160C) J head residues (C), or Smc1(K201C) and Smc3(K198C) coiled coil residues (D) was performed in vivo using BMOE. Cell extracts were labelled with HaloTag-TMR ligand. Smc-HaloTag species were separated by SDS-PAGE and quantified by in-gel fluorescence. Percentage of cross-link efficiency is given.

According to the modelled J head Sc Smc structure, Smc1S161C and Smc3K160C are predicted to be 9.1 Å apart (Fig 2B) and BMOE induced moderately efficient cross-linking dependent on the presence of both cysteines (Figs 2C and S1D). Interestingly, the incidence of cross-linking was doubled by Smc3K38I or Smc3S1127R mutations that greatly reduce ATP-dependent head engagement. These data confirm that Smc1 and Smc3 heads can also adopt the J conformation, which is presumably incompatible with E detected by BMOE induced cross-linking between Smc1N1192C and Smc3R1222C. If so, the high incidence of E Smc1/3 head engagement may be at the expense of J complexes. Thus, Smc3K38I and Smc3S1127R may increase the J state by reducing the E state.

BMOE induced efficient cross-linking between Smc3K198C and Smc1K201C dependent on both cysteine substitutions (Figs 2D and S1E). Surprisingly, this crosslinking between Smc1 and Smc3 coiled coils was unaffected by either Smc3K38I or Smc3S1127R mutations and only modestly reduced by Smc3E1155Q (Fig. 2D), raising the possibility that the interaction detected by cross-linking Smc3K198C to Smc1K201C might occur with E as well as J heads.

### The state of cohesin’s hinge and coiled coils when its ATPase heads are engaged

To address the state of cohesin’s hinge when its heads are in E and J states and when its coiled coils are juxtaposed, we measured the products of cross linking when cysteine pairs were present at two interfaces. The incidence of crosslinking between Smc1K639C and SmcE570C within cohesin’s hinge was 24% while that between Smc1N1192C and Smc3R1222C within the E interface was 45%. Cross-linking both interfaces simultaneously produces a product that migrates slightly faster than that produced by hinge cross-linking alone (Fig. 3A). TEV-engineered cleavage confirmed that this species was due to simultaneous cross-linking of hinge and E interfaces from same Smc1/3 heterodimer (Fig. S2A). Importantly, the incidence of the doubly crosslinked product was 13%, suggesting that the probability of crosslinking at both interfaces is close to the product of the probabilities of cross-linking at each site (0.45 × 0.24 = 0.11). Hinge crosslinking therefore is unaffected by cross-linking associated with the E state (Fig. 3A) and vice versa. This implies that ATP-driven head engagement does not cause any appreciable opening of the hinge interface, at least not in a manner that would hinder cross-linking between Smc1K639C and SmcE570C. The incidence of cross-linking at the interface between Smc1/3 coiled coils (using Smc3K198C and Smc1K201C) together that of the hinge was 12% (Fig. 3B), which was twice as high as expected (6%) if the two events were unconnected, suggesting that crosslinking one interface may actually increase the chances at the other.

**Figure 3.**
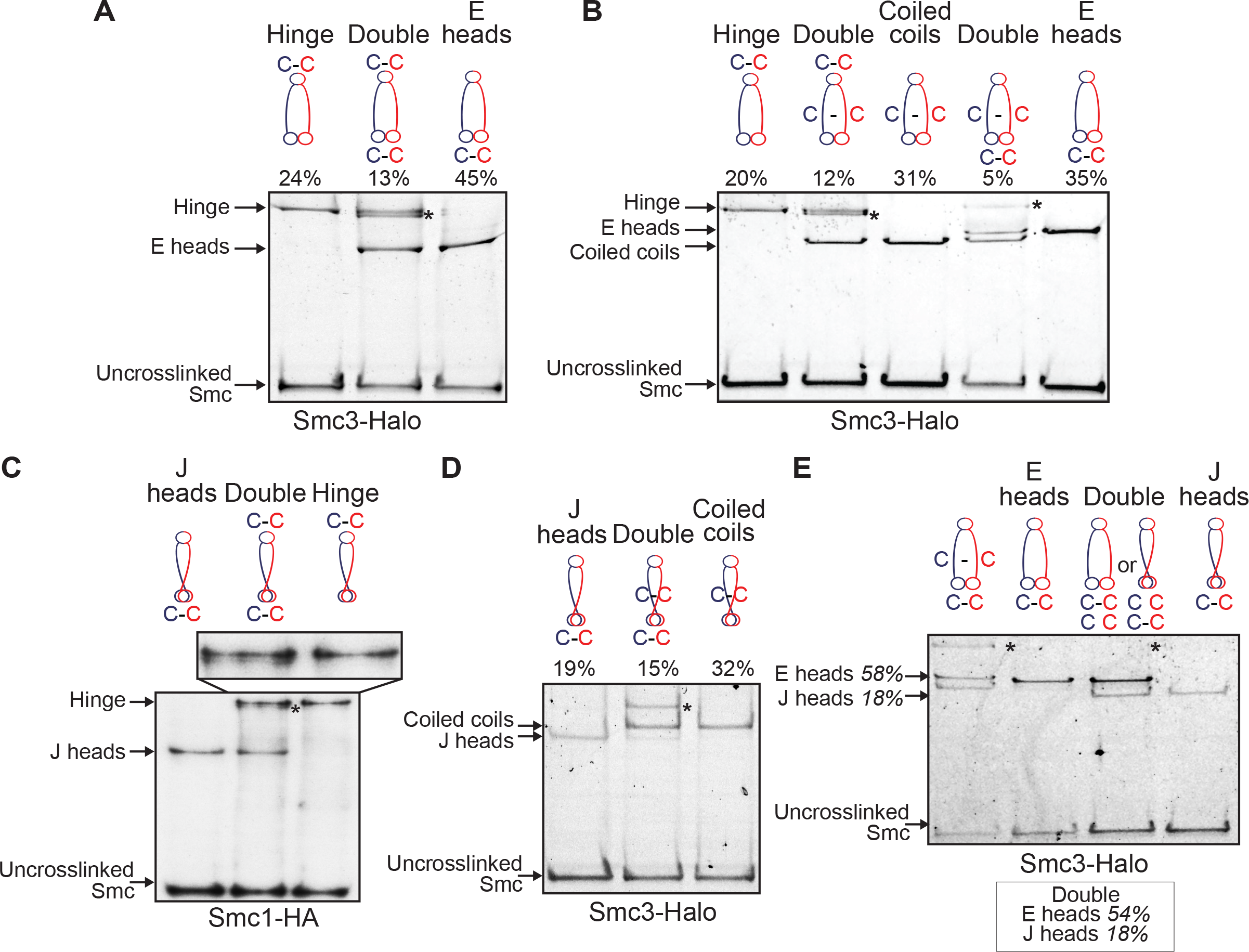
S compartments: Coiled coils, E and J head interactions in cohesin heterodimers. (A) E state of Smc heads in heterodimers. Smc1 and Smc3-HaloTag proteins containing hinge and/or E heads cysteine pairs were cross-linked in vivo using BMOE. Complexes were immunoprecipitated on Scc1-PK6, labelled with TMR ligand, separated by SDS-PAGE and quantified by in-gel fluorescence. Percentage of cross-linking efficiency is given. Asterisk shows the location of the double crosslink. (B) E Smc heads and coiled coils interactions in heterodimers. Smc1 and Smc3-HaloTag proteins containing hinge, E heads and/or coiled coils cysteine pairs were analysed as in (A). (C) J state of Smc heads in heterodimers. Smc1-HA, Smc3 and Scc1-PK proteins containing hinge and/or J heads cysteine pairs were were crosslinked in vivo, immunoprecipitated on Scc1-PK6, separated by SDS-PAGE and analysed by western blot. (D) J heads interactions with Smc coiled coils. Smc1 and Smc3-HaloTag proteins containing coiled coils and/or alternative J heads cysteine pairs were analysed as in (A). (E) E and J states of Smc heads are mutually exclusive. Smc1 and Smc3-HaloTag proteins containing E heads and/or J heads cysteine pairs were analysed as in (A). Percentage of the double cross-link efficiency is shown in box. Left lane: double crosslink of E heads with coiled coils shown for size indication.

In contrast, the incidence of simultaneous cross-linking at coiled coil and E interfaces (5%) was lower than expected (11%) were the two events unconnected, suggesting that crosslinking at one site is associated with a reduction in the incidence of cross-linking at the other (Fig. 3B, see also S2B and C). It is nevertheless striking that the Smc1/3 coils associated with E heads can in fact be cross-linked, albeit with a lower than expected probability. In other words, the E state does not seem to eliminate the possibility of crosslinking Smc1/3 coiled coils, at least between Smc3K198C and Smc1K201C. At first glance, this finding is difficult to reconcile with the notion that close juxtaposition of Smc coiled coils in the vicinity of their joints drives formation of the J state (Diebold-Durand et al., 2017). However, we note that the same double cross-linking experiment was never performed with B. subtilis proteins and we cannot therefore conclude that cohesin is substantially different from Bs Smcs in this regard.

In summary, the incidence of double cross-linking suggests that Smc1/3 heterodimers with engaged heads are also connected via their hinges, as are complexes whose coiled coils are juxtaposed. In contrast, head engagement appears to be associated with a lower than expected probability of coiled coil juxtaposition.

### ATP-dependent head engagement (E) and signature motif juxtaposition (J) are distinct states

We were able to detect dimers arising from simultaneous cross-linking of hinge and J head interfaces by western-blot (Fig. 3C) but not by in-gel fluorescence, presumably because they co-migrate with those cross-linked merely at their hinges in the presence of a Halo-tag on Smc3 (Fig. S2D). However, our finding that hinge cross-linking is accompanied by a modest reduction in the incidence of dimers cross-linked solely between Smc1S161C and Smc3K160C indicates that double crosslinking does indeed occur. Likewise, due to the proximity of their cysteine insertions within the N-terminal domains of Smc1 and Smc3, it would not have been possible to distinguish the mobility of dimers cross-linked simultaneously between coiled coil (Smc3K198C and Smc1K201C) and J head (Smc1S161C and Smc3K160C) interfaces. Nevertheless, our finding that mutations like Smc3K38I and Smc3S1127R, which largely eliminate E but increase J, modestly increase Smc3K198C/Smc1K201C crosslinking (Fig. 2D) suggests that coiled coils are indeed juxtaposed in the J conformation.

To answer this question more definitively, we used an alternative cysteine pair within the coiled coil joint region (Smc3E202C and Smc1R1031C, Hu & Than, personal communication). Unlike Smc1K201C, R1031C is C-terminal and crosslinking with Smc3E202C altered the mobility of dimers created by cross-linking Smc1S161C and Smc3K160C (Fig. 3D and S2B). Moreover, the incidence of simultaneous cross-linking at this new coiled coil interface together with the J interface (15%) was more than twice that expected (6%) were the two events unconnected. In other words, crosslinking one interface increased the chances of the other, unlike the situation with E heads (Fig. 3B). It would therefore appear that Smc1 and Smc3 coiled coils in the vicinity of the joint are more frequently associated when heads are in the J state than the E state.

Because of the primary sequence proximity of coiled coil (Smc3K198C and Smc1K201C) and J (Smc1S161C and Smc3K160C) cysteine pairs, the migration of dimers cross-linked simultaneously at coiled coil and E interfaces indicates where dimers cross-linked simultaneously at J and E interfaces would migrate. However, BMOE treatment of cells containing cysteine pairs at both J and E interfaces revealed no dimers with the migration expected of simultaneous crosslinking (Fig. 3E and S2C), implying that cross-linking at J and E interfaces are mutually exclusive. According to the E and J models, Smc1S161/Smc3K160 and Smc1N1192 /Smc3R1222 are 25.4 Å and 43.5 Å apart respectively (Fig. 1C and 2B), distances that would not permit BMOE-induced cross-linking. Thus, cross-linking between Smc1S161C/Smc3K160C and Smc1N1192C /Smc3R1222C pairs reveal distinct J and E states.

### Identification of K compartments associated with both J and E states

To address whether the state of ATPase heads affects their association with Scc1’s N- and C-terminal domains (NScc1 and CScc1), we measured cross-linking between cysteine pairs at the Smc3 neck/NScc1 or the Smc1 head/CScc1 interfaces in cells that also contained cysteine pairs at E or J interfaces. In all four combinations, the fraction of simultaneous cross-linking was similar or equal to the product of the fractions of molecules crosslinked at individual interfaces (Figs. 4A-D). Because NScc1 and CScc1 are usually bound to Smc3 and Smc1 simultaneously (Gligoris et al., 2014), we conclude that both E and J states give rise to K compartments defined by Smc heads that associate simultaneously with both ends of Scc1 as well as with themselves. Double cross-linking experiments demonstrated that juxtaposition of Smc1 and Smc3 coiled coils also takes place when Scc1 is connected to their head and necks respectively (Fig. S2E and F).

**Figure 4.**
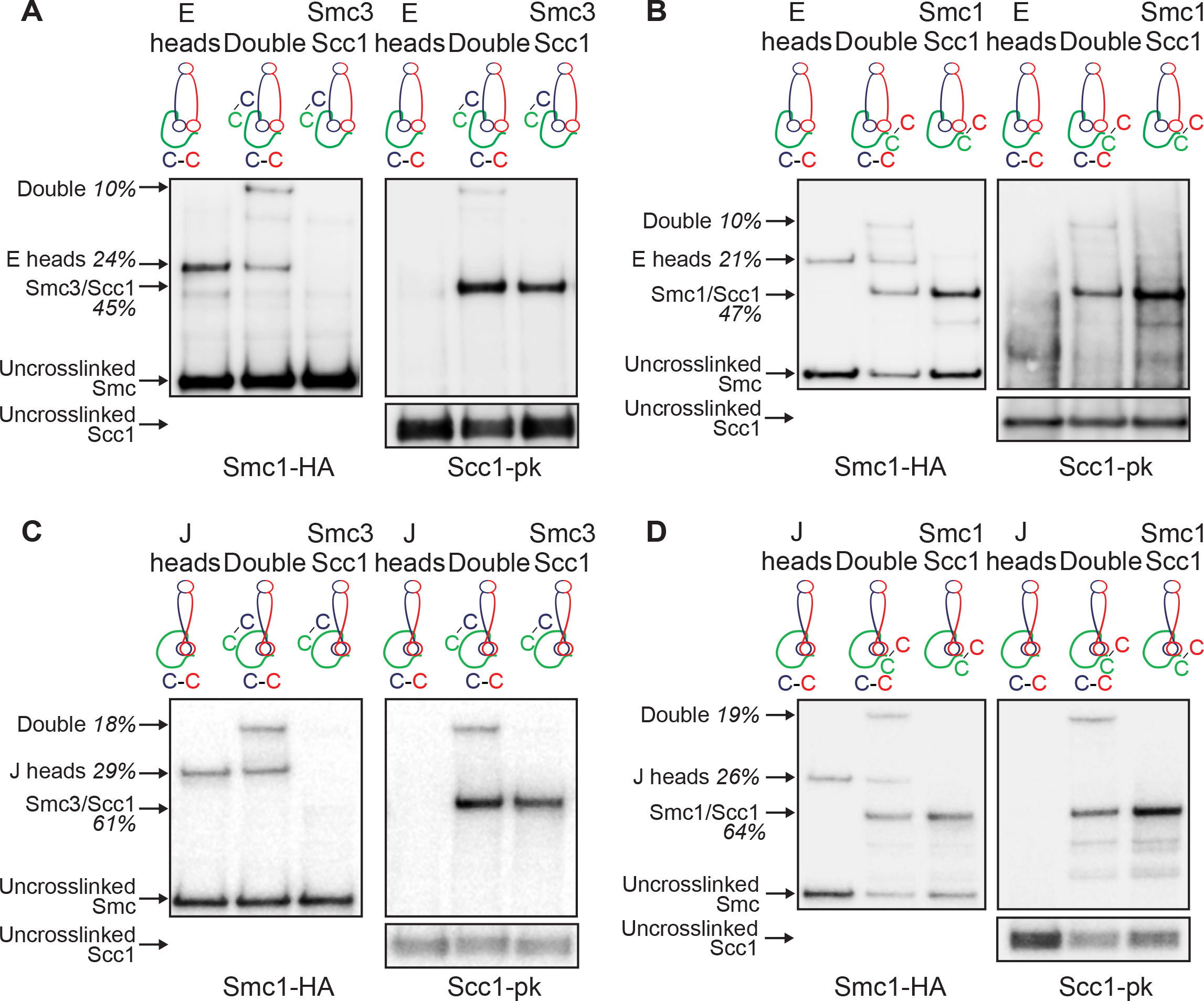
K compartments: E and J head interactions with Scc1. (A) E heads interact with Scc1 at Smc3-Scc1 interface. Smc1-HA, Smc3 and Scc1-PK proteins containing E heads or Smc3-Scc1 interface cysteine were crosslinked in vivo, immunoprecipitated on Scc1-PK6, separated by SDS-PAGE and semi-quantified by western blot. Percentage of cross-link efficiency is given. (B) E heads interact with Scc1 at Smc1-Scc1 interface. Smc1-HA, Smc3 and Scc1-PK proteins containing E heads or Smc1-Scc1 interface cysteine were analyzed as in (A). (C) J heads interact with Scc1 at Smc3-Scc1 interface. Smc1-HA, Smc3 and Scc1-PK proteins containing J heads or Smc3-Scc1 interface cysteine were analyzed as in (A). (D) J heads interact with Scc1 at Smc1-Scc1 interface. Smc1-HA, Smc3 and Scc1-PK proteins containing J heads or Smc1-Scc1 interface cysteine were analyzed as in (A).

### E and J states occur throughout the cell cycle

The incidence of E heads detected by Smc1N1192C /Smc3R1222C crosslinking was only modestly lower in cells arrested in G2/M by nocodazole than in cells arrested in late G1 by expression of non-degradable Sic1 (Fig. 5A and S3A). It was also unaffected by inactivation of Scc2, Hos1, Wapl, or Smc1D1164E (Fig. 5C and S3D), a mutation that like *wpl1*Δ abolishes releasing activity (Beckouet et al., 2016; Camdere et al., 2015; Elbatsh et al., 2016), or by Pds5 depletion (Fig. 5D and S3E). Remarkably, it was also unaffected by Scc1 depletion (Figs. 5B, S3B, and S3C), implying that ATP can induce head engagement within Smc1/3 heterodimers. J heads detected by Smc1S161C/Smc3K160C crosslinking were more frequent in G2/M than in late G1 (Fig. 5A and S3F), and reduced in cells depleted either of Pds5 (Figs. 5E, S3G, and S3I) or Eco1 (Figs. 5F, S3H and S3I).

**Figure 5.**
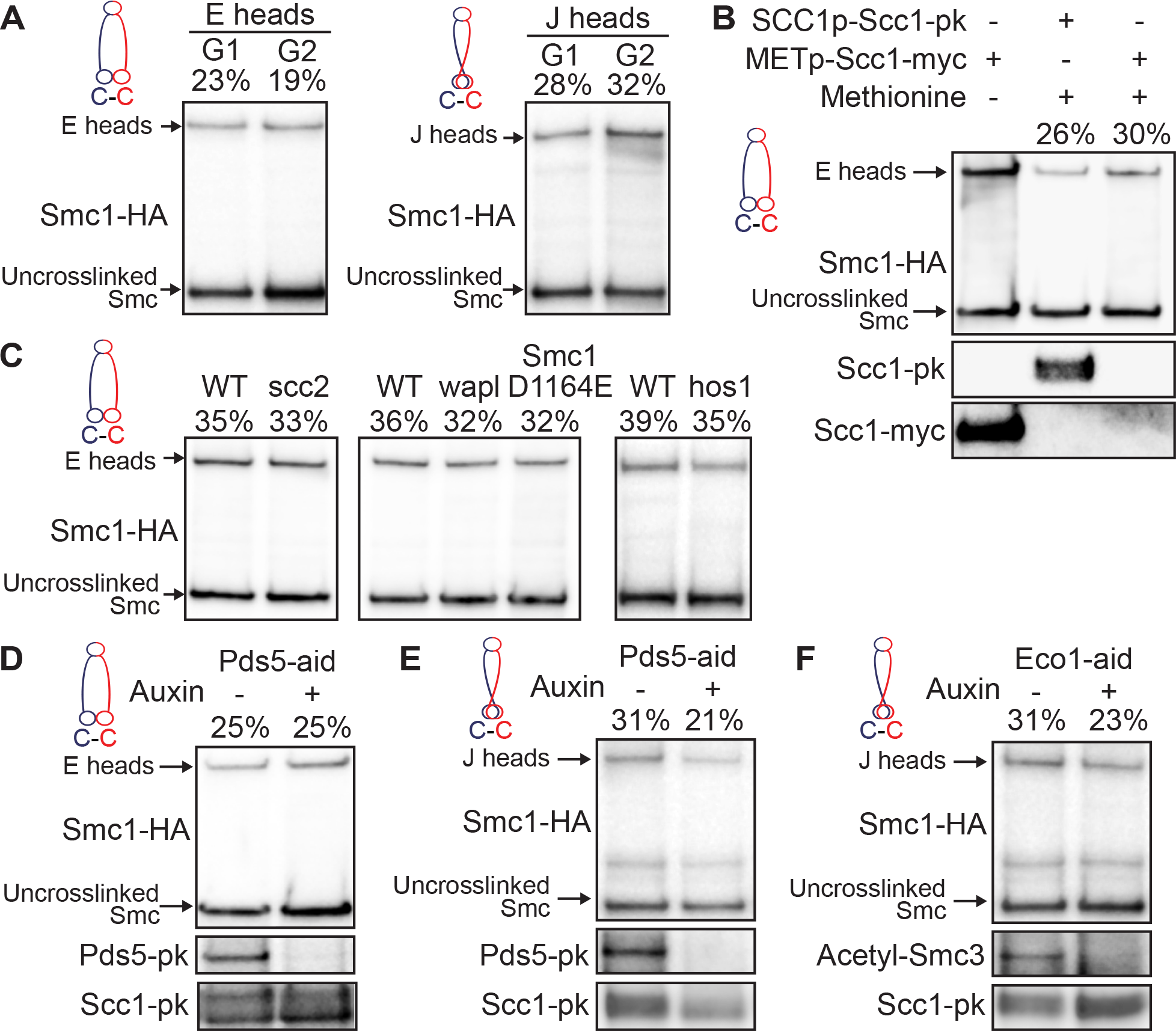
Head engagement and juxtaposition occur throughout the cell cycle. (A) E and J heads over the cell cycle. Smc1-HA and Smc3 proteins containing E or J heads cysteine pairs were cross-linked in G1 (Gal-Sic1) or G2/M (nocodazole) arrested cells using BMOE. Complexes were immunoprecipitated on Scc1-PK, separated by SDS-PAGE and analysed by western blot. Percentage of cross-link efficiency is given. (B) E heads upon absence of Scc1. Cells were pheromone-arrested in early G1 prior to Scc1-myc transcription repression using an inducible methionine promoter and release into nocodazole. Smc1-HA and Smc3 proteins containing E heads cysteine pairs were cross-linked in vivo using BMOE. Protein extracts were analyzed by SDS-PAGE and western blot. Left lane: cycling cells expressing Scc1-myc under control of methionine promoter were analyzed in parallel. (C) E heads upon absence of functional cohesin loader Scc2 or releasing activity. Smc1-HA and Smc3 proteins containing E heads cysteine pairs were cross-linked in vivo using BMOE. Complexes were analyzed as in (A). Left panel: WT and scc2-45 strains were arrested in G1 with alpha factor at 25°C and released into nocodazole at 37°C. Middle panel: cycling WT, *wpl* deleted and Smc1(D1164E, N1192C)-HA strains. Right panel: cycling WT and *hos1* deleted strains. (D) E heads in the absence of Pds5. Pds5-aid strain was arrested in G1 with alpha factor, supplemented with auxin for 1 h prior to release into nocodazole-containing media supplemented with auxin. In vivo cross-linked proteins were analyzed as in (A). (E-F) J heads in the absence of Pds5 (H) or Eco1 (I). WT, Pds5-aid or Eco1-aid strains were analyzed as in (D).

### Sister DNAs are entrapped within J-K compartments

Armed with cysteine pairs within E or J interfaces that can be crosslinked simultaneously either with those at hinge or Smc-kleisin interfaces, we used gel electrophoresis to measure entrapment of DNAs within all four types of S and K compartments. Cycling cells from strains containing a variety of cysteine pairs were treated with BMOE and circular minichromosome DNAs associated with cohesin were separated by electrophoresis following denaturation with SDS. In addition to nicked and supercoiled monomeric DNAs, cells containing cysteine pairs at hinge and both Smc-kleisin interfaces contain monomeric supercoiled DNAs whose migration is modestly retarded due to their catenation by a single cohesin ring (catenated monomers -CMs) as wells as monomeric supercoiled DNAs that migrate as dimers due to their co-entrapment within a single cohesin ring (catenated dimers -CDs)(Fig. 6A). Importantly, neither CMs nor CDs were observed when hinge cysteine pairs were combined with those at either E or J interfaces, implying that DNAs are rarely, if ever, entrapped within either type of S compartment (Fig. 6AB and S4A). In contrast, both CMs and CDs were observed when cysteine pairs at both Smc-kleisin interfaces were combined with the J pair (Fig. 6B and S4B). CMs were also detected, albeit at a low level, when cysteine pairs at both Smc-kleisin interfaces were instead combined with those at the E interface. CDs on the other hand, were never detected under these conditions (Fig. 6A and S4A). Thus, while both types of K compartments can entrap individual DNAs, only J-K compartments can entrap sisters.

**Figure 6.**
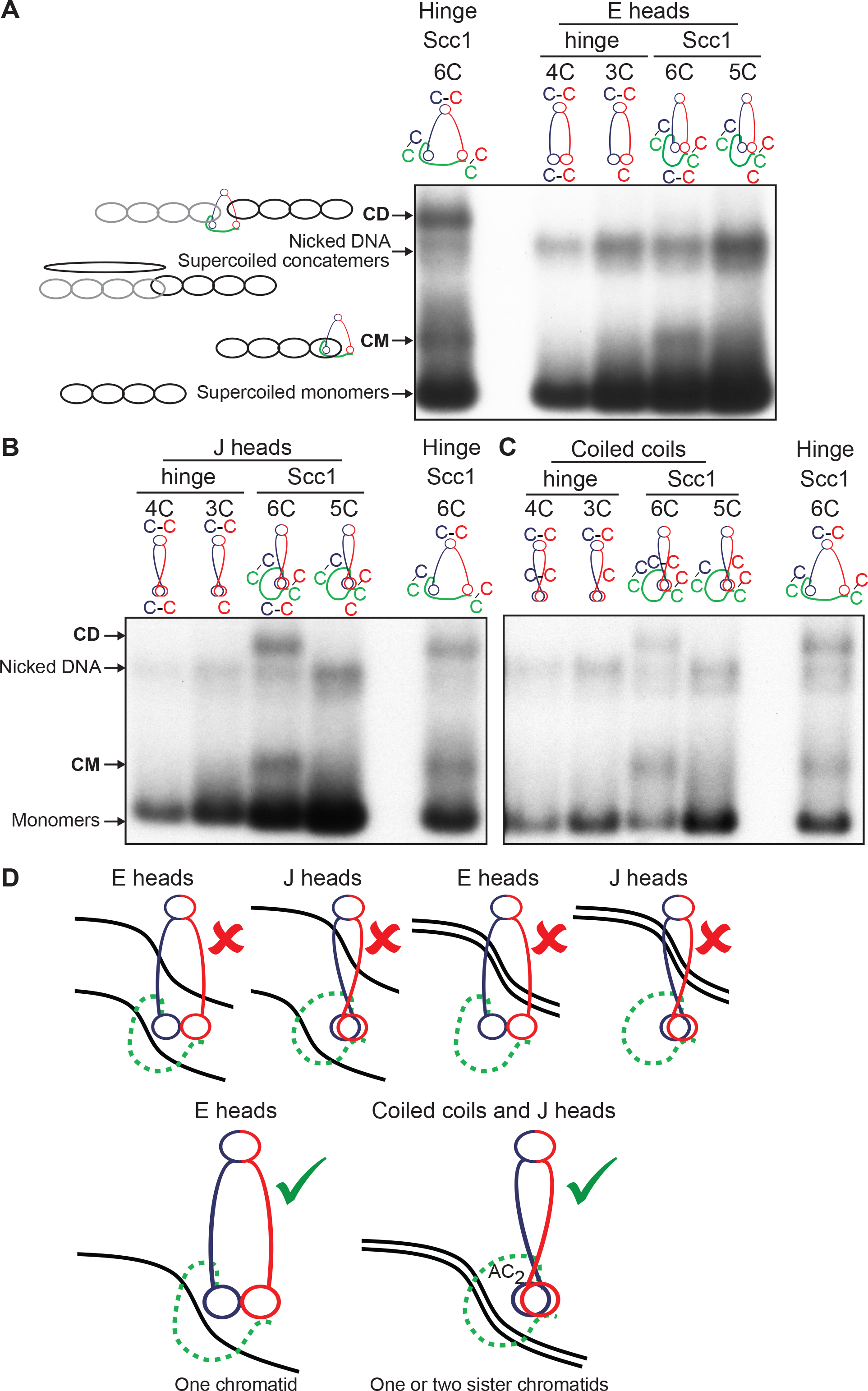
CMs and CDs in core K compartments with E and J heads. (A) CMs and CDs in exponentially growing strains containing cysteines in the hinge, E heads or Smc1/Scc1/Smc3 interfaces. Strains with cysteine pairs at interfaces (4C or 6C) and strains lacking just one of the cysteines (3C or 5C), carrying a 2.3 kb circular minichromosome, were treated with BMOE. DNAs associated with cohesion immunoprecipitates (Scc1-PK) were denatured with SDS and separated by agarose gel electrophoresis. Southern blotting reveals supercoiled monomers and nicked and supercoiled concatemers along with two forms of DNA unique to 6C cells, termed CMs and CDs (See Gligoris et al., Science 2014 for details). (B) CMs and CDs in exponentially growing strains containing cysteines in the hinge, J heads or Smc1/Scc1/Smc3 interfaces. (C) CMs and CDs in exponentially growing strains containing cysteines in the hinge, coiled coils or Smc1/Scc1/Smc3 interfaces. (D) Schematic representation of DNA topological association with cohesin compartments.

As predicted by work on bacterial Smcs showing that the coiled coils associated with J kleisin-compartments are juxtaposed, CMs and CDs were also detected when cysteines at both Smc-kleisin interfaces were combined with cysteines at the Smc1/3 coiled coil interface, close to the joint region (Fig. 6C). Importantly, neither form of entrapment was detected when cysteines at the Smc1/3 coiled coil interface were combined with cysteines at the hinge (Fig. 6C), which is consistent with the lack of DNA entrapment within S compartments.

Our failure to detect entrapment of DNAs inside S compartments is inconsistent with the prevailing view that cohesin holds sister chromatids together by entrapping DNAs between the hinges and heads of its Smc1/3 subunits. Our findings suggest that DNAs are instead entrapped in the K compartment created by the binding of Scc1’s N- and C-terminal domain to Smc3 and Smc1 heads whose signature motifs are juxtaposed in the absence of ATP. In other words, Scc1’s association with Smc1/3 heterodimers creates the K compartment that entraps DNAs. Scc1 does not merely strengthen an S compartment created by the association of Smc1 and Smc3 via their hinges and heads. Interactions between Smc1/3 coiled coils may contribute to the exclusion of DNAs from S compartments, except fleetingly perhaps during loading or translocation reactions. Because cross-linking traps interactions between proteins, it cannot reveal information on its dynamics. Thus, the observation that DNAs are entrapped by J-K compartments does not exclude the possibility that transient dissociation of Smc heads gives rise to open SK rings that would also maintain entrapment. Indeed, the notion that J-K compartments are in a dynamic equilibrium with open SK rings is the simplest way of explaining how cleavage of Smc3 coiled coils is sufficient to release cohesin from DNA (Gruber et al., 2003; Li et al., 2017; Murayama and Uhlmann, 2015).

### J heads are a feature of sister chromatid cohesion throughout the genome

A clear limitation of the CM/CD assay is that it only measures the state of minichromosome DNAs. Even in this case, it is only revealing about states of association between DNA and cohesin that involve their topological catenation. To address whether cohesion throughout the genome is mediated by J head rather than E head cohesin, we used antibodies specific for acetylated and non-acetylated Smc3 to detect cross-linking between J and E interfaces. Though the frequencies of cross-linking of acetylated and non-acetylated Smc3 hinges to Smc1 were similar (Fig. 7A and S4C), E-specific crosslinking was far less frequent with acetylated Smc3 (3%) than with non-acetylated Smc3 (41%) (Fig. 7A and S4C). Strikingly, the opposite was true for J-specific crosslinking. Thus, 7% of non-acetylated and 37% of acetylated Smc3 were crosslinked at the J interface (Fig. 7A and S4C). Because acetylation is associated with cohesion establishment during S phase, these data suggest that cohesin complexes engaged in holding sisters together throughout the genome are much more likely to be in the J than the E state. The data raise the possibility that cohesion itself might directly prevent ATP-dependent head engagement. In addition to Smc heads being more likely to be in the J than the E state, acetylated chromosomal cohesin appears to be have a tighter or more frequent association with Pds5 (Chan et al., 2013). Thus, calibrated ChiP-seq revealed that inactivation of Eco1 reduced Pds5’s association throughout the genome compared to that of Scc1, an effect that was even more pronounced in *wpl1* mutants (Figs. 7B, 7C, 7D, S4D, S4E, S5, S6). This observation is consistent with in vivo FRAP measurements suggesting that acetylation reduces Pds5 turnover on chromosomes (Chan et al., 2013).

**Figure 7.**
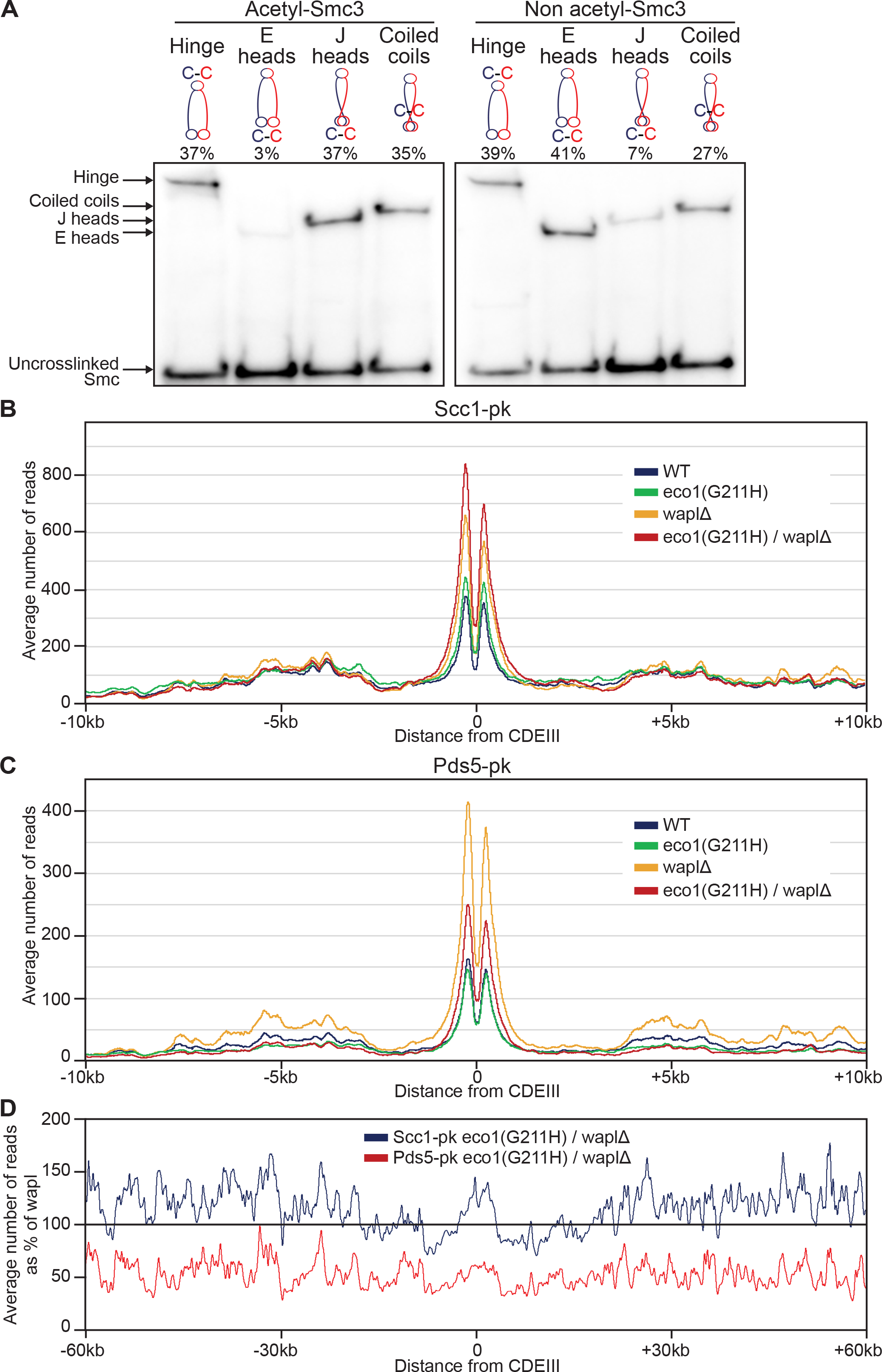
Acetylation mediated control of both J head and Pds5 chromatin association. (A) Smc3 acetylation of heterodimers with E and J heads. Smc1-HA and Smc3 proteins containing hinge, E heads, J heads or coiled coils cysteine pairs were cross-linked in vivo with BMOE. Complexes were immunoprecipitated on Scc1-PK, separated by SDS-PAGE and analyzed by western blot against acetylated-Smc3 or non-acetylated Smc3. Percentage of the cross-link signal is given for each antibody. (B-C) Average calibrated ChIP-seq profiles of Scc1-PK (B) and Pds5-PK (C) in the presence or absence of Wapl and/or functional Eco1. Cells were arrested in G2 using nocodazole at restrictive temperature after release from pheromone arrest at permissive temperature. (D) Averaged calibrated ChIP-seq profiles 60 kb either side of CDEIII plotted as a percentage of the average number of reads of Scc1-PK and Pds5-PK obtained for *wapl*Δ cells in (B) and (C) respectively.

## Discussion

Sister chromatid cohesion is a feature of chromosome segregation that is universal among eukaryotes and a property that distinguishes them from bacteria. An important clue regarding the mechanism was the finding that the Scc1, Smc1, and Smc3 subunits of the cohesin complex responsible bind each other in a pairwise manner to create a huge tripartite ring whose cleavage by separase triggers the dissolution of cohesion at anaphase. This raised the possibility that cohesin holds sister DNAs together using a topological principle, namely co-entrapment of sister DNAs inside the tripartite SK ring formed by the binding of Scc1’s N- and C-terminal domains to the necks and heads of Smc3 and Smc1 that are themselves associated via their hinges (Haering et al., 2008; Haering et al., 2002). Though previous thiol-specific cross-linking studies have confirmed the entrapment of sister minichromosome DNAs inside such rings (Gligoris et al., 2014), they have not hitherto taken into account another key feature of Smc/kleisin complexes, namely that the ATPase heads at the vertices of V-shaped Smc1/3 heterodimers can themselves interact, thereby dividing the ring into two compartments, a S compartment created by association of Smc1 hinges and heads with equivalent domains within Smc3 and a K compartment created by juxtaposition of Smc1 and Smc3 heads that are also associated with the C- and N-terminal domains of Scc1 (Arumugam et al., 2003). Crosslinking studies using a series of novel cysteine pairs described here show that cohesin’s Smc1 and Smc3 ATPase heads in fact associate in two very different modes, the canonical one that sandwiches a pair of ATP molecules between engaged heads (E) and another in which the heads are rotated and translocated in a fashion that juxtaposes their signature motifs in the absence of ATP (J). The transition between these two states driven by the binding and hydrolysis of ATP, first discovered in bacteria (Diebold-Durand et al., 2017), may be a universal feature of Smc/kleisin complexes (Buermann et al., 2018). There are accordingly two types of S and K compartments, those associated with J or E heads.

The previous finding that sister DNAs are entrapped by rings cross-linked at hinge and both Smc-kleisin interfaces is consistent with several scenarios (Fig. 6D). DNAs could be entrapped in open SK rings but never in ones whose heads are in either E or J mode. Both DNAs could be entrapped either in J-S or in E-S compartments. Both could be entrapped either in J-K or in E-K-compartments. Lastly, one DNA could be entrapped in an E-S or in a J-S compartment while the other in the associated K compartment. Our new cross-linking studies rule out all but one of these scenarios and imply that both sister DNAs are in fact frequently entrapped in J-K but not E-K compartments. The lack of entrapment either of individual or sister DNAs in either type of S compartment excludes the possibility that both DNAs are entrapped in a S compartment or that one DNA is entrapped within a S compartment and the second within its associated K compartment. DNA entrapment in J-K compartments is also found in B.subtilis (Vazquez Nunez et al.) and this mode of association may therefore be universal.

The finding that acetylation of Smc3 during S phase is accompanied by a pronounced increase in J but decrease in E suggests that entrapment of sister DNAs within J-K compartments may be a feature of cohesion throughout the genome and does not merely apply to small circular minichromosomes. The observation that sister DNAs are entrapped in J-K compartments refines our view of cohesion while the failure to observe entrapment exclusively by open SK rings, S compartments, E-K compartments, or entrapment of one DNA in an S and its sister in a K compartment exclude most previously proposed scenarios (Huber et al., 2016; Li et al., 2017; Murayama et al., 2018; Murayama and Uhlmann, 2015). It is nevertheless important to point out that detection of sister DNA entrapment by J-K compartments does not exclude the possibility that J-K compartments are in dynamic equilibrium with open SK rings. Though sister DNAs were never observed in E-K compartments, individual DNAs were, albeit infrequently. An explanation for this finding is that ATP-driven head engagement necessary for E-K compartment entrapment cannot occur when sister DNAs are present.

To explain why cleavage of Smc3’s coiled coil by separase alleviates the retardation of sister chromatid disjunction caused by *hos1* mutations, it was suggested that sister DNAs are normally entrapped in an E-S compartment and that the Smc3 acetylation that persists in *hos1* mutations blocks the head disengagement necessary for DNA escape via a gate created by kleisin cleavage (Li et al., 2017). Our failure to observe stable entrapment of DNAs in either J-S or E-S compartments is inconsistent with this hypothesis and implies that there must be an alternative interpretation for the *hos1* effect. We suggest that Smc3 de-acetylation is instead required to facilitate the escape of DNAs from J-K compartments upon Scc1 cleavage, possibly by weakening an association of Pds5 with Scc3 or Smc1 heads that would otherwise hinder escape.

Because formation of J-K compartments is likely to require ATP hydrolysis, the notion of cohesion being mediated by entrapment of sister DNAs within J-K compartments is hard to reconcile with the proposal that ATP hydrolysis is unnecessary for building sister chromatid cohesion. The argument that hydrolysis is not required is based on the behaviour of Smc1D1164E mutants that can load onto chromosomes and build cohesion despite being defective in ATP hydrolysis (Camdere et al., 2018). Though these mutations may reduce ATP hydrolysis, we suggest that their viability in fact depends on residual ATPase activity. Cohesin containing Smc1E1158Q Smc3E1155Q, which is completely defective in ATP hydrolysis (Petela et al., 2018), cannot load correctly onto chromosomes let alone build cohesion (Arumugam et al., 2006; Hu et al., 2011; Hu et al., 2015).

One reason why DNAs are not entrapped in J-S compartments is that the coiled coils associated with the Smc1 and Smc3 heads are juxtaposed throughout their length, as suggested for bacterial Smc proteins and eukaryotic cohesin (Buermann et al., 2018; Diebold-Durand et al., 2017; Kulemzina et al., 2016). Lack of DNA entrapment in E-S compartments is more surprising given that it is widely assumed (by analogy with Rad50) that ATP-driven head engagement creates a DNA binding groove that would be situated within the E-S compartment (Liu et al., 2016; Rojowska et al., 2014; Schuler and Sjogren, 2016; Seifert et al., 2016). If such a groove is also a feature of the E state in cohesin, then DNA binding at this site must be infrequent but could nevertheless be an important, albeit transient, event during the process of DNA entrapment. Alternatively, DNAs might lie within the groove as part of a loop (i.e. passage of DNA through the hole twice, once in one direction and once in the opposite), which would not result in S compartment entrapment as measured in our assay. Though our experiments shed important insight into the eventual location of sister DNAs within the cohesin ring, future experiments will be required to explore the series of events that create this state, in particular whether a fleeting entrapment within E-S compartments is involved, how DNAs enter cohesin rings, and how sister DNAs enter the same J-K compartment. Lastly, the topology of cohesin’s association with DNA when extruding loops also remains to be explored.

## Supporting information

## Acknowledgments

We are grateful to Katsuhiko Shirahige for supplying antibodies, to Bin Hu and Thane Than for sharing with us unpublished coiled coil crosslinking data and to Stephan Gruber and Roberto Vazquez Nunez for sharing with us unpublished data. We thank previous and current members of the Nasmyth and Brockdorff groups for valuable discussions and technical assistance, Frederick Beckouet, Martin Houlard and Jean Metson. This work was funded by the Cancer Research UK (12386 and 26747 to K.A.N.), Wellcome Trust (107935/Z/15/Z to K.A.N.) and European Research Council (294401 to K.A.N).

## Author contributions

C.C. and K.N. wrote the manuscript. C.C. designed and conducted experiments. R.J. and T.v.O. conducted experiments. J.C.S. aided with initial technical procedures.

## Supplementary Figures and table

**Figure S1. Related to Figure 1 and 2**

(A) Multiple sequence alignment indicating the lack of conservation of Smc residues which were mutated to cysteines.

(B) Cells containing indicated cysteine residues were grown in YPD at different temperatures (25°C, 30°C and 37°C). Coiled coils, E and J heads cysteine pairs do not rescue lethality of eco1(G211H) temperature sensitive mutant at 37°C.

(C) In vivo cysteine cross-linking of Smc1(N1192C)-HA6 and Smc3(R1222C)-PK6 proteins depends on both cysteine residues and BMOE. Crosslinking reaction was performed in vivo with BMOE. Cell protein extracts were separated by SDS-PAGE and analysed by western blot.

(D) In vivo cysteine cross-linking of Smc1(S161C)-HA6 and Smc3(K160C)-PK6 proteins depends on both cysteine residues and BMOE. Crosslinking reaction was performed in vivo with BMOE. Cell protein extracts were separated by SDS-PAGE and analysed by western blot.

(E) In vivo cysteine cross-linking of Smc1(K201C)-myc9 and Smc3(K198C)-PK6 proteins depends on both cysteine residues and BMOE. Crosslinking reaction was performed in vivo with BMOE. Cell protein extracts were separated by SDS-PAGE and analysed by western blot.

**Figure S2. Related to Figure 3 and 4**

(A) Smc1(K639C)-HA6 and Smc3(E570C, 968/969flagTEVx3)-PK6 or Smc1(K639C, N1192C)-HA6 and Smc3(E570C, 968/969flagTEVx3, R1222C)-PK6 proteins containing hinge and/or E heads cysteine pairs were cross-linked in vivo using BMOE. Complexes were immunoprecipitated on Smc3-PK6, cut by recombinant Tev protease, separated by SDS-PAGE and western blot.

(B) Smc1 and Smc3-HaloTag proteins containing E heads, alternative J heads and/or coiled coils cysteine pairs were cross-linked in vivo with BMOE. Complexes were immunoprecipitated on Scc1-PK6, labelled with TMR ligand, separated by SDS-PAGE and quantified using in-gel fluorescence. Percentage of cross-link efficiency is given. Asterisk shows the location of the double crosslink.

(C) Smc1 and Smc3-HaloTag proteins containing E heads, J heads and/or coiled coils cysteine pairs were analyzed as in (B).

(D) Left panel: Smc1-HA, Smc3 and Scc1-PK proteins containing hinge and/or J heads cysteine pairs were were crosslinked in vivo, immunoprecipitated on Scc1-PK6, separated by SDS-PAGE and analysed by western blot. Right panel: Smc1 and Smc3-HaloTag proteins containing hinge and/or J heads cysteine pairs were analysed as in (B). Percentage of the double cross-link efficiency is indicated in box.

(E-F) Coiled coils interactions in trimers. Smc1-HA, Smc3 and Scc1-pk6 proteins containing coiled coils cysteine pairs combined with cysteine pairs at the Smc3-Scc1 (C) or Smc1-Scc1 (D) interface were cross-linked in vivo using BMOE. Complexes were immunoprecipitated on Scc1-PK6, separated by SDS-PAGE and semi-quantified by western blot. Percentage of cross-link efficiency is given.

**Figure S3. Related to Figure 5**

(A) FACS profiles for the experiment described in Figure 5A.

(B) Strains containing the Scc1 gene under the control of the WT Scc1 or Met3-repressible promoter as described in Figure 5B were grown in synthetic media lacking methionine or YPD.

(C) FACS profiles for the experiment described in Figure 5B.

(D) FACS profiles for the experiment described in Figure 5C.

(E) FACS profiles for the experiment described in Figure 5D.

(F) FACS profiles for the experiment described in Figure 5A.

(G) FACS profiles for the experiment described in Figure 5E.

(H) FACS profiles for the experiment described in Figure 5F.

(I) Strains containing Pds5-AID or Eco-AID do not grow on YPD + 5mM Auxin

**Figure S4. Related to Figures 6 and 7**

(A) CMs and CDs in exponentially growing strains containing cysteines in the hinge, E heads and/or Smc1/Scc1/Smc3 interfaces.

(B) CMs and CDs in exponentially growing strains containing cysteines in the hinge, J heads and/or Smc1/Scc1/Smc3 interfaces.

(C) Smc1-HA and Smc3 proteins containing hinge, E heads, J heads or coiled coils cysteine pairs were cross-linked in vivo using BMOE and analyzed as in Figure 7A.

(D) FACS profiles for the experiment described in Figure 7B.

(E) FACS profiles for the experiment described in Figure 7C.

**Figure S5. Related to Figure 7**

(A) Calibrated ChIP-seq profiles of Scc1-PK6 in the absence of Wapl and in the presence of WT Eco1 (blue) or the absence of both Wapl and functional Eco1 (orange). Sixteen yeast chromosomes profiles are centred on the CDEIII element.

(B) Calibrated ChIP-seq profiles of Scc1-PK6 on chromosomes I and IV in the absence of Wapl, plus/minus functional Eco1.

**Figure S6. Related to Figure 7**

(A) Calibrated ChIP-seq profiles of Pds5-PK6 in the absence of Wapl and presence of WT Eco1 (blue) or the absence of both Wapl and functional Eco1 (orange). Sixteen yeast chromosomes profiles are centred on the CDEIII element.

(B) Calibrated ChIP-seq profiles of Pds5-PK6 on chromosomes I and IV in the absence of Wapl, plus/minus functional Eco1.

**Table S1.**
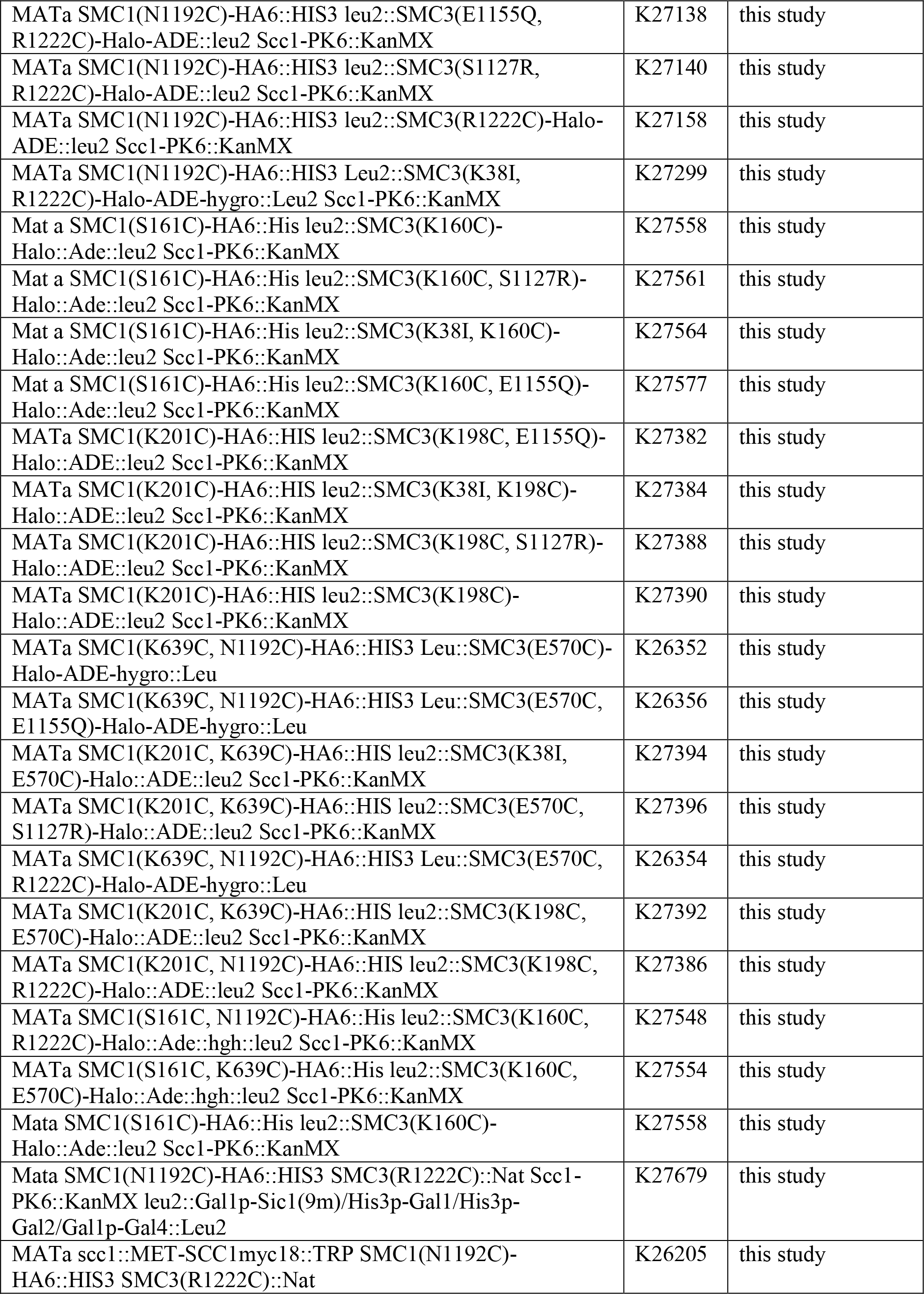
Strains used in this study. All strains used in this study are derived from K699 (S.Cerevisiae W303).

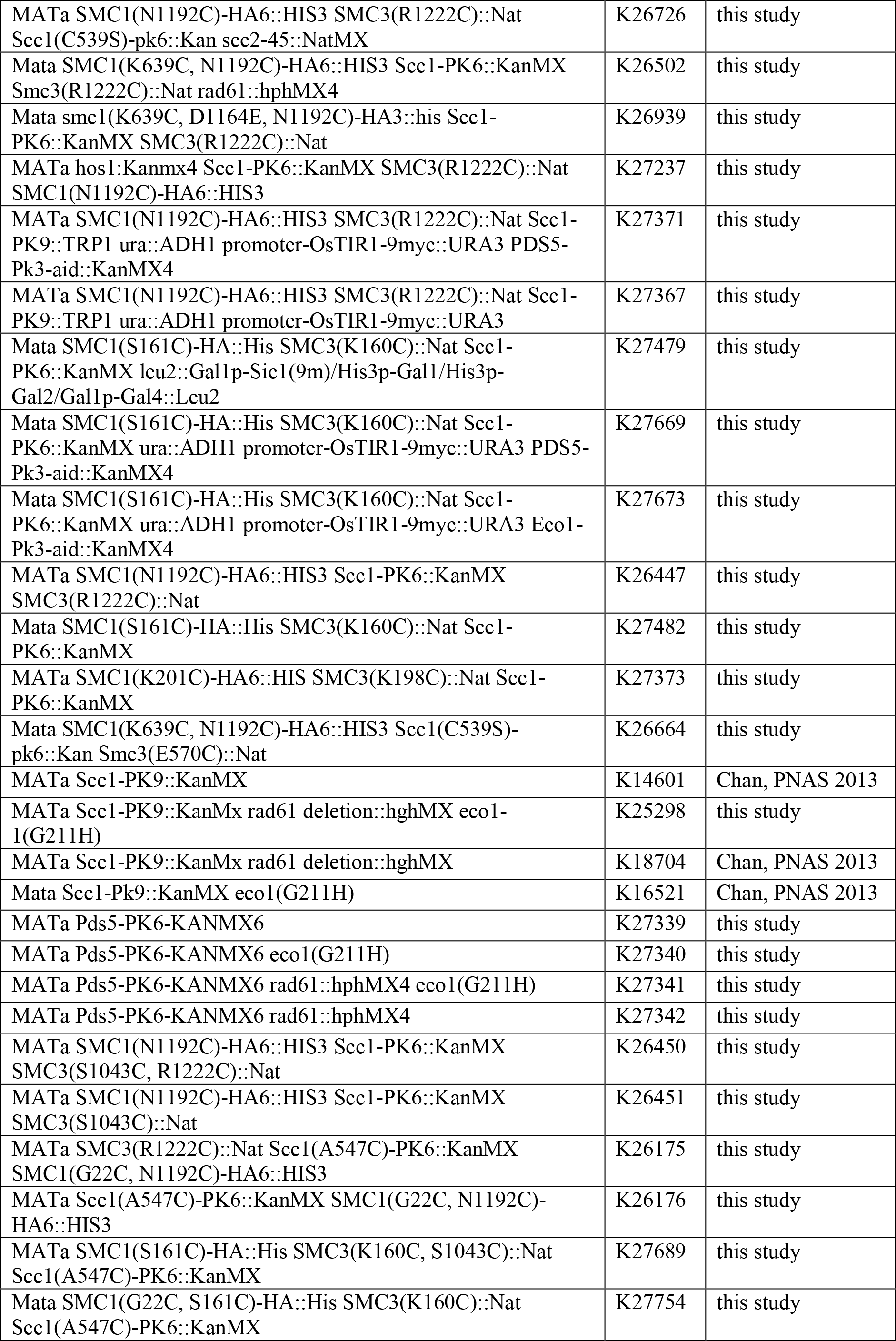

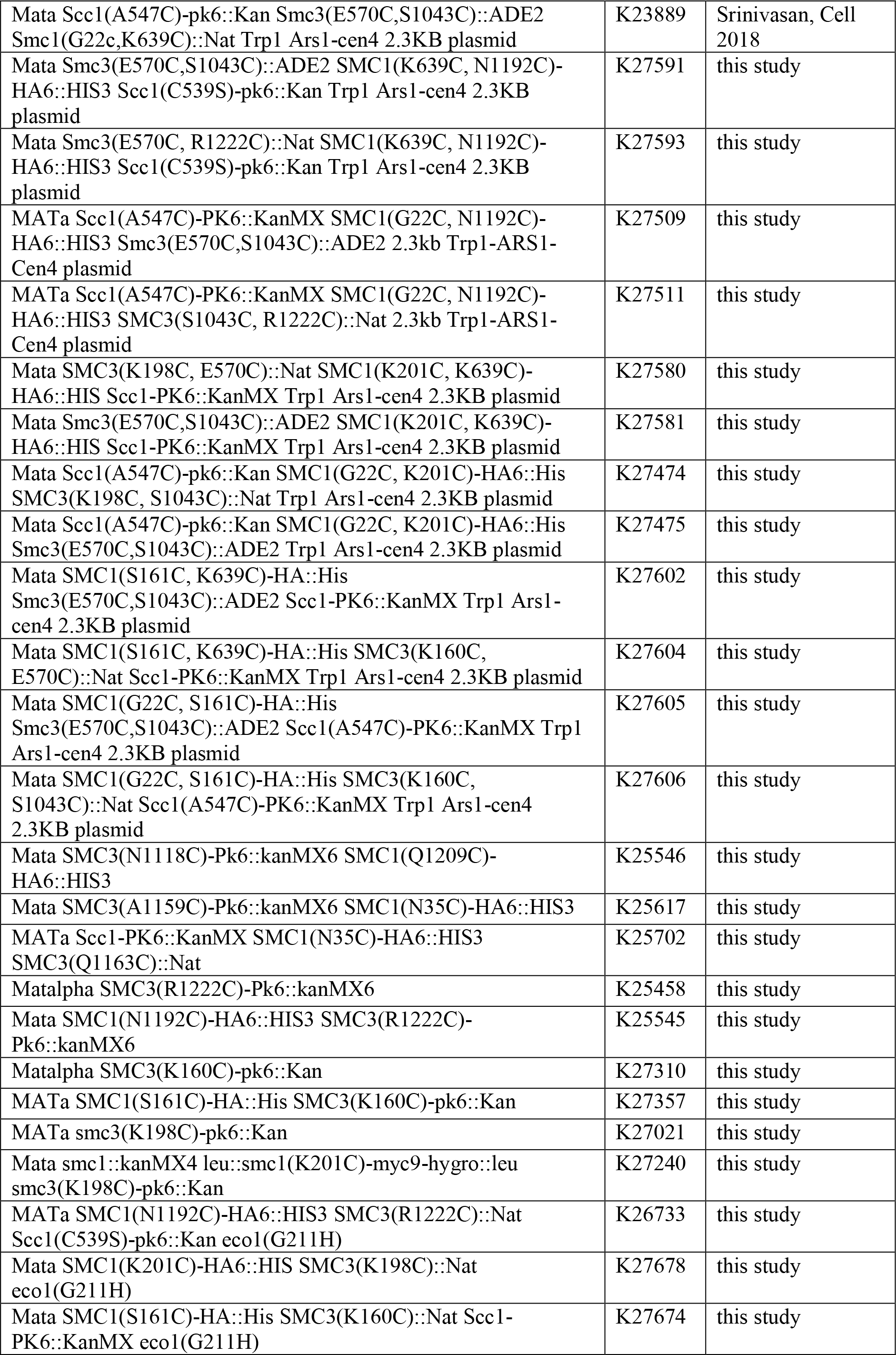

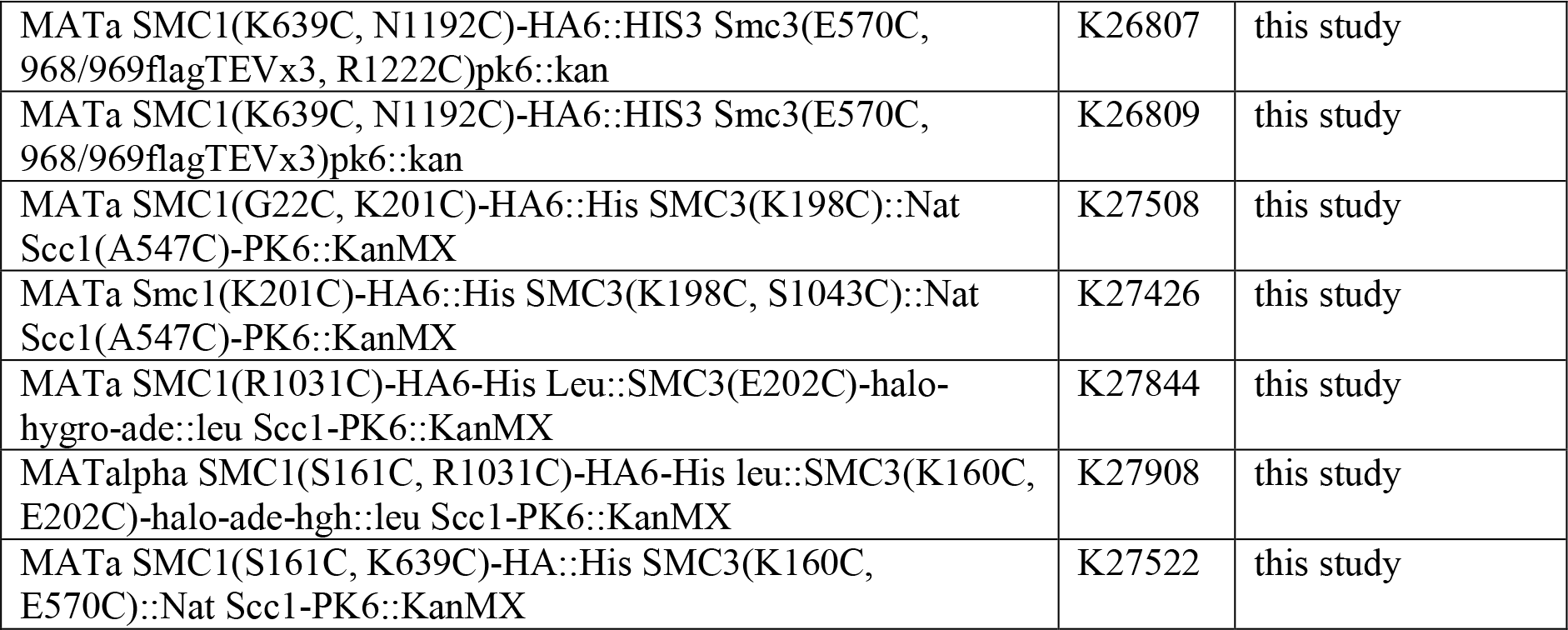

## Material and method

### Yeast cell culture

All strains are derivatives of W303 (K699). Strain numbers and relevant genotypes of the strains used are listed in the supplementary Table S1. Cells were cultured at 25C in YEP medium with 2% glucose unless stated otherwise. To arrest the cells in G1, a-factor was added to a final concentration of 2 mg/L, every 30 min for 2.5 h. Cells were released from G1 arrest by filtration wherein cells were captured on 1.2 mm filtration paper (Whatman GE Healthcare), washed with 1 L YEPD and resuspended in the appropriate fresh media. To inactivate temperature sensitive alleles, fresh media was pre-warmed prior to filtration (Aquatron, Infors HT). To arrest cells in G2, nocodazole (Sigma) was added to the fresh media to a final concentration of 10 mg/mL and cells were incubated until the synchronization was achieved (> 95% large-budded cells). Cells were arrested in late G1 by galactose-induced overexpression of a non-degradable mutant of the Sic1 protein (mutation of 9 residues phosphorylated by Cdk1). To achieve this, cells were grown in YEP supplemented with 2% raffinose and arrested in G1 as described above. 1 h before release from G1 arrest, galactose was added to 2% of the final concentration. Cells were released into YEPD as described above, and incubated for 60 min at 25C. For auxin induced degradation of proteins, cells were arrested in G1 as above and 1 h prior to release auxin (indole-3-acetic acid sodium salt; Sigma) was added to a final concentration of 5 mM. Cells were released from G1 arrest into YEPD medium containing 5 mM auxin and 10 mg/mL nocodazole. To produce cells deficient of Scc1, the gene was placed under the MET3-repressible promoter. Liquid cultures were grown in minimal media supplemented with 2% glucose and 1%-MET dropout solution overnight, diluted to OD600 = 0.2 and allowed to grow to OD600 = 0.4. Cells were then collected by filtration as described above, resuspended in YPD supplemented with 8mM methionine and arrested in G1. Once arrested, the cells were collected by filtration, washed with YPD in the presence of 8mM methionine and released into the same media.

### In vivo chemical crosslinking and protein detection

Strains were grown in YEPD at 25C to OD600nm = 0.5-0.6. 15 OD units were washed in ice-cold PBS and re-suspended in 1 mL icecold PBS. The suspensions were then split into 2 × 300 uL and 12.5uL BMOE (stock: 125 mM in DMSO, 5 mM final) or DMSO was added for 6 min on ice. Cells were washed with 2 × 2 mL ice-cold PBS containing 5 mM DTT, resuspended in 750 uL lysis buffer (25 mM HEPES pH 8.0, 50 mM KCl, 50 mM MgSO4, 10 mM trisodium citrate, 25 mM sodium sulfite, 0.25% triton-X, freshly supplemented with Roche Complete Protease Inhibitors (2X) and PMSF (1 mM), lysed in a FastPrep-24 (MP Biomedicals) for 3 × 1 min at 6.5 m/s with 750uL of acid-washed glass beads (425-600 mm, Sigma) and lysates cleared (5 min, 12 kg). Protein concentrations were adjusted after Bradford assay and cohesin immuno-precipitated using either anti-PK antibody (AbD Serotec) or anti-HA antibody (Roche 3F10, 1 h, 4C) and protein G dynabeads (1 h, 4C, with rotation) in presence of Halo-Tag TMR ligand (Promega). Beads were washed with 2 × 1mL lysis buffer, resuspended in 50uL 2× sample buffer, incubated at 95C for 5 min and the supernatant loaded onto a 3%-8% Tris-acetate gradient gel (Life Technologies). Gels were scanned with an FLA7000 scanner (Fuji) and processed for Western blotting. The proteins were then transferred onto Immun-Blot PVDF using Trans-blot Turbo transfer packs for the Trans-blot Turbo system (Bio-Rad). For visualization the membrane was incubated with appropriate antibodies (Mouse anti-PK, AbD Serotec; Rat anti-HA antibody, Roche 3F10; Mouse anti-Myc, Millipore 4A6; Mouse anti-flag, Sigma-Aldrich M2; Mouse monoclonal anti acetylated Smc3, H2 gift from Katsuhiro Shirahige; Mouse monoclonal anti unacetylated Smc3, Bio6 described in Chan et al. PNAS 2013; Goat anti-Rat HRP, Millipore; Sheep anti-Mouse HRP, GE Healthcare) and with Immobilon Western Chemiluminescent HRP substrate (Millipore) before detection using an ODYSSEY Fc Imaging System (LI-COR). Intensity of each cross-linked band was calculated as a percentage of total signal intensity of the lane.

### Minichromosome IP

Strains containing a 2.3 kb circular minichromosome harboring the TRP1 gene were grown overnight in ‒TRP medium at 25C and sub-cultured in YEPD medium for exponential growth (OD600nm = 0.6). 15 OD units were washed in ice-cold PBS and processed for in vivo crosslinking as described above with the following modification: after cohesin immuno-precipitation protein G dynabeads were washed with 2 × 1 mL lysis buffer, resuspended in 30 uL 1% SDS with DNA loading dye, incubated at 65C for 4 min and the supernatant run on a 0.8% agarose gel containing ethidium bromide (1.4 V/cm, 22h, 4C). After Southern blotting using alkaline transfer, bands were detected using a 32-P labeled TRP1 probe.

### Multiple sequence alignment

Multiple sequence alignments were created using Clustal Omega (Sievers et al., 2011). The following sequences were included: Homo sapiens, Mus musculus, Drosophila melanogaster, Saccharomyces cerevisiae, Schizosaccharomyces pombe, Pyrococcus furiosus, Pyrococcus yayanosii, Bacillus subtilis.

### Calibrated ChIP-sequencing

Cells were grown exponentially to OD600 = 0.5 and the required cell cycle stage where necessary. 15 OD600nm units of S. cerevisiae cells were then mixed with 3 OD600nm units of C. glabrata to a total volume of 45 mL and fixed with 4 mL of fixative (50 mM Tris-HCl, pH 8.0; 100 mM NaCl; 0.5 mM EGTA; 1 mM EDTA; 30% (v/v) formaldehyde) for 30 min at room temperature (RT) with rotation. The fixative was quenched with 2 mL of 2.5 M glycine (RT, 5 min with rotation). The cells were then harvested by centrifugation at 3,500 rpm for 3 min and washed with ice-cold PBS. The cells were then resuspended in 300 mL of ChIP lysis buffer (50 mM HEPESKOH, pH 8.0; 140 mM NaCl; 1 mM EDTA; 1% (v/v) Triton X-100; 0.1% (w/v) sodium deoxycholate; 1 mM PMSF; 2X Complete protease inhibitor cocktail (Roche)) and an equal amount of acid-washed glass beads (425-600 mm, Sigma) added before cells were lysed using a FastPrep_-24 benchtop homogenizer (M.P. Biomedicals) at 4C (3 × 60 s at 6.5 m/s or until > 90% of the cells were lysed as confirmed by microscopy). The soluble fraction was isolated by centrifugation at 2,000 rpm for 3 min then sonicated using a bioruptor (Diagenode) for 30 min in bursts of 30 s on/30 s off at high level in a 4C water bath to produce sheared chromatin with a size range of 200-1,000 bp. After sonication the samples were centrifuged at 13,200 rpm at 4C for 20 min and the supernatant was transferred into 700 uL of ChIP lysis buffer. 30 uL of protein G Dynabeads (Invitrogen) were added and the samples were pre-cleared for 1 h at 4C. 80 uL of the supernatant was removed (termed ‘whole cell extract [WCE] sample’) and 5 mg of antibody (anti-PK (Bio-Rad) or anti-HA (Roche)) was added to the remaining supernatant which was then incubated overnight at 4C. 50 uL of protein G Dynabeads were then added and incubated at 4C for 2 h before washing 2× with ChIP lysis buffer, 3× with high salt ChIP lysis buffer (50mMHEPES-KOH, pH 8.0; 500 mM NaCl; 1 mM EDTA; 1% (v/v) Triton X-100; 0.1% (w/v) sodium deoxycholate; 1 mM PMSF), 2× with ChIP wash buffer (10 mM Tris-HCl, pH 8.0; 0.25MLiCl; 0.5% NP-40; 0.5% sodium deoxycholate; 1mM EDTA; 1 mMPMSF) and 1× with TE pH7.5. The immunoprecipitated chromatin was then eluted by incubation in 120 uL TES buffer (50 mM Tris-HCl, pH 8.0; 10 mM EDTA; 1% SDS) for 15 min at 65C and the collected supernatant termed ‘IP sample’. The WCE samples were mixed with 40 uL of TES3 buffer (50 mM Tris-HCl, pH 8.0; 10 mM EDTA; 3% SDS) and all samples were de-crosslinked by incubation at 65C overnight. RNA was degraded by incubation with 2 uL RNase A (10 mg/mL; Roche) for 1 h at 37C and protein was removed by incubation with 10 uL of proteinase K (18 mg/mL; Roche) for 2 h at 65C. DNA was purified using ChIP DNA Clean and Concentrator kit (Zymo Research).

### Preparation of sequencing libraries

Sequencing libraries were prepared using NEBNext Fast DNA Library Prep Set for Ion Torrent Kit (New England Biolabs) according to the manufacturer’s instructions. Briefly, 10-100ng of fragmented DNA was converted to blunt ends by end repair before ligation of the Ion Xpress Barcode Adaptors. Fragments of 300bp were then selected using E-Gel SizeSelect 2% agarose gels (Life Technologies) and amplified with 6-8 PCR cycles. The DNA concentration was then determined by qPCR using Ion Torrent DNA standards (Kapa Biosystems) as a reference. 12-16 libraries with different barcodes could then be pooled together to a final concentration of 350pM and loaded onto the Ion PI V3 Chip (Life Technologies) using the Ion Chef (Life Technologies). Sequencing was then completed on the Ion Torrent Proton (Life Technologies), typically producing 6-10 million reads per library with an average read length of 190bp.

### Data analysis, alignment, and production of BigWigs

Unless otherwise specified, data analysis was performed on the Galaxy platform (Giardine et al., 2005). Quality of reads was assessed using FastQC (Galaxy tool version 1.0.0) and trimmed as required using ‘trim sequences’ (Galaxy tool version 1.0.0). Generally, this involved removing the first 10 bases and any bases after the 200th, but trimming more or fewer bases may be required to ensure the removal of kmers and that the per-base sequence content is equal across the reads. Reads shorter than 50bp were removed using Filter FASTQ (Galaxy tool version 1.0.0, minimum size: 50, maximum size: 0, minimum quality: 0, maximum quality: 0, maximum number of bases allowed outside of quality range: 0, paired end data: false) and the remaining reads aligned to the necessary genome(s) using Bowtie2 (Galaxy tool version 0.2) with the default (‒sensitive) parameters (mate paired: single-end, write unaligned reads to separate file: true, reference genome: SacCer3 or CanGla, specify read group: false, parameter settings: full parameter list, type of alignment: end to end, preset option: sensitive, disallow gaps within n-positions of read: 4, trim n-bases from 50 of each read: 0, number of reads to be aligned: 0, strand directions: both, log mapping time: false). To generate alignments of reads that uniquely align to the S. cerevisiae genome, the reads were first aligned to the C. glabrata (CBS138, genolevures) genome with the unaligned reads saved as a separate file. These reads that could not be aligned to the C. glabrata genome were then aligned to the S. cerevisiae (sacCer3, SGD) genome and the resulting BAM file converted to BigWig (Galaxy tool version 0.1.0) for visualization. Similarly, this process was done with the order of genomes reversed to produce alignments of reads that uniquely align to C. glabrata.

### Visualization of ChIP-seq profiles

The resulting BigWigs were visualized using the IGB browser (Nicol et al., 2009). To normalize the data to show quantitative ChIP signal the track was multiplied by the samples’ occupancy ratio (OR) and normalized to 1 million reads using the graph multiply function. In order to calculate the average occupancy at each base pair up to 60 kb around all 16 centromeres, the BAM file that contains reads uniquely aligning to S. cerevisiae was separated into files for each chromosome using ‘Filter SAM or BAM’ (Galaxy tool version 1.1.0). A pileup of each chromosome was then obtained using samtools mpileup (Galaxy tool version 0.0.1) (source for reference list: locally cached, reference genome: SacCer3, genotype likelihood computation: false, advanced options: basic). These files were then amended using our own script (chr_position.py) to assign all unrepresented genome positions a value of 0. Each pileup was then filtered using another in-house script (filter.py) to obtain the number of reads at each base pair within up to 60 kb intervals either side of the centromeric CDEIII elements of each chromosome. The number of reads covering each site as one successively moves away from these CDEIII elements could then be averaged across all 16 chromosomes and calibrated by multiplying by the samples’ OR and normalizing to 1 million reads.

## DATA AND SOFTWARE AVAILABILITY

### Scripts

All scripts written for this analysis method are available to download from https://github.com/naomipetela/nasmythlab-ngs.

### Calibrated ChIP-seq data

The GEO accession number for the calibrated ChIP-seq data (raw and analyzed) reported in this paper is: GSE120138.

