## Supplementary figures and images for "The topology of DNA entrapment by cohesin rings"

### Supplementary file 1

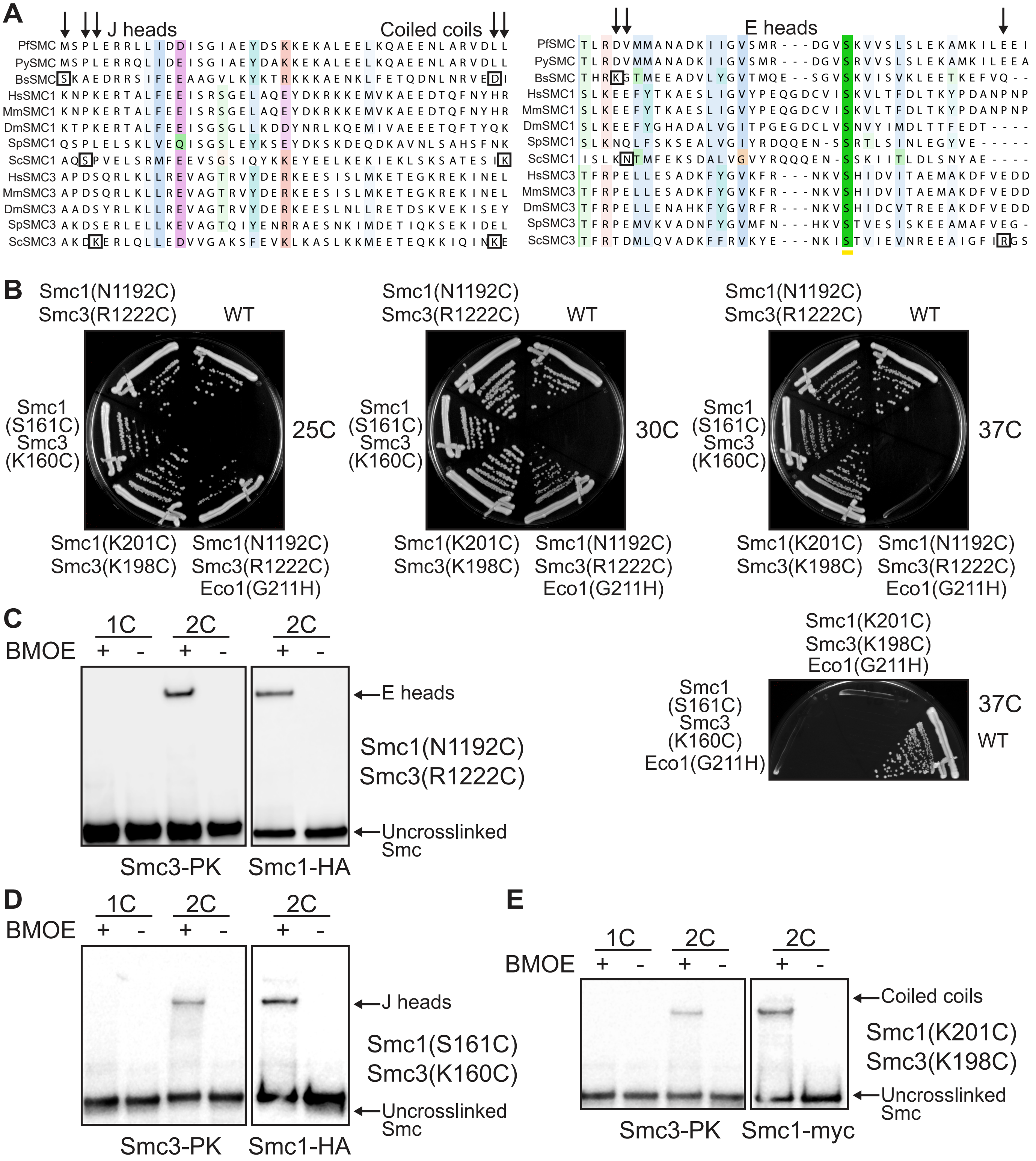

### Supplementary file 2

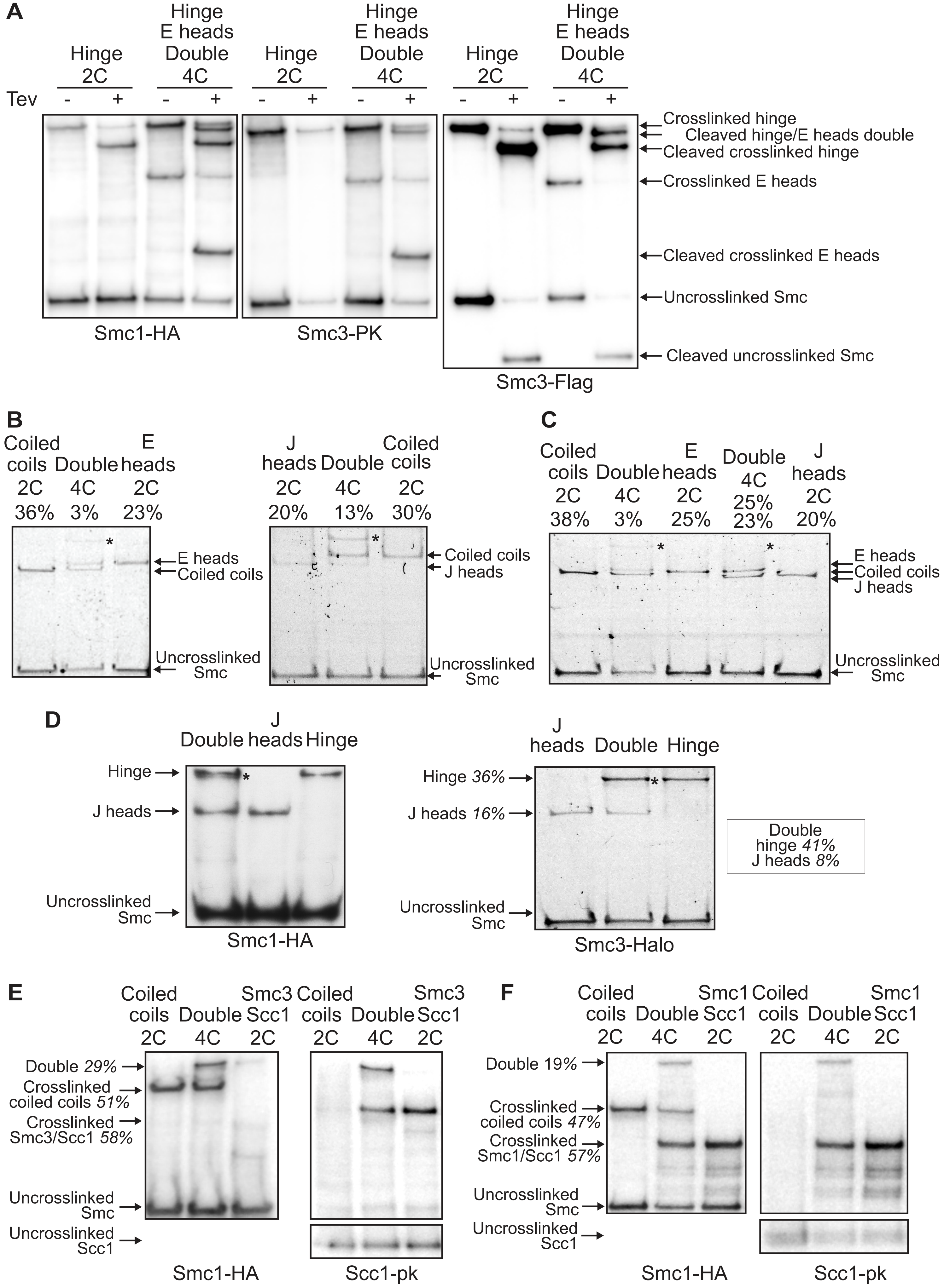

### Supplementary file 3

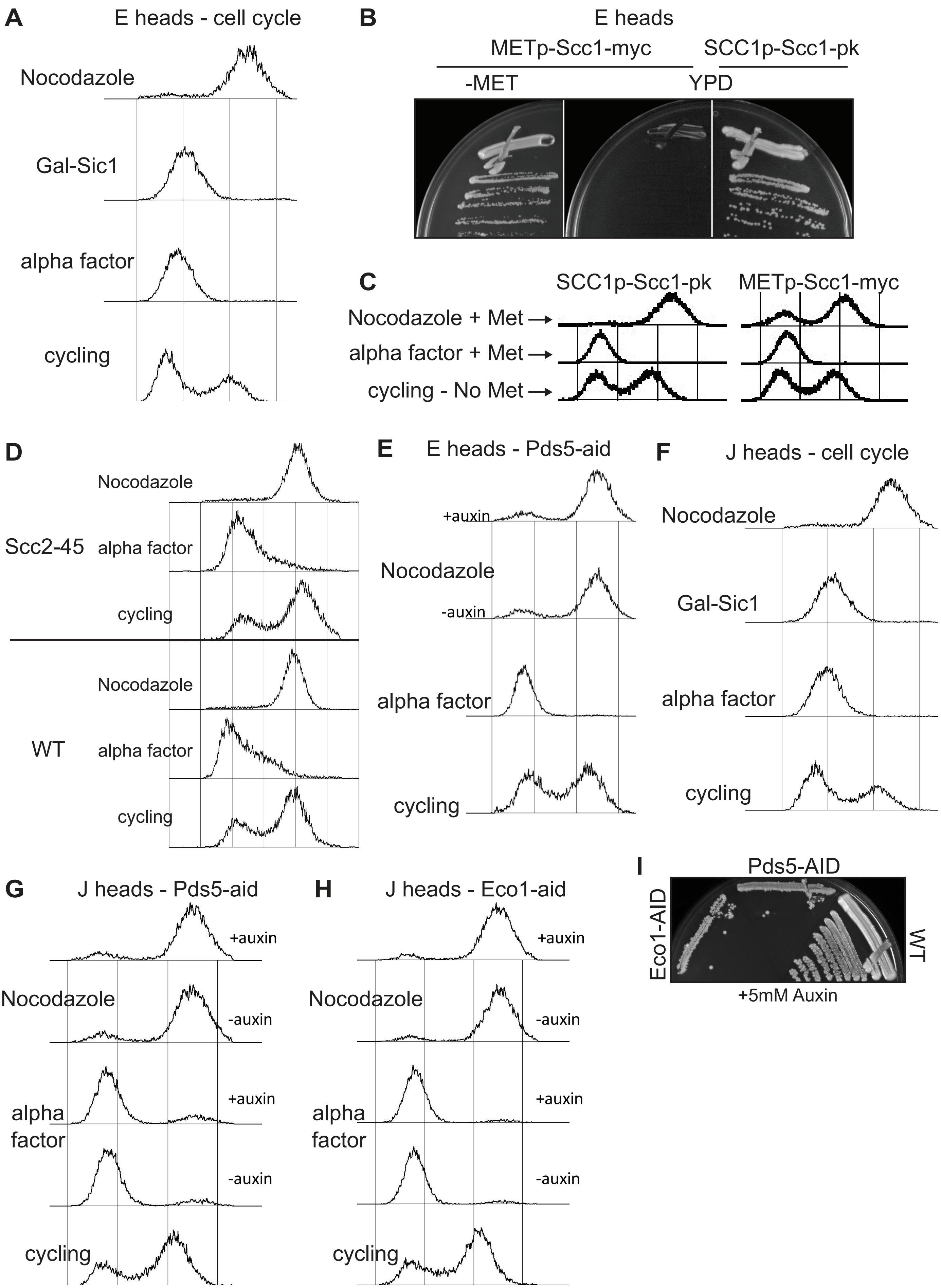

### Supplementary file 4

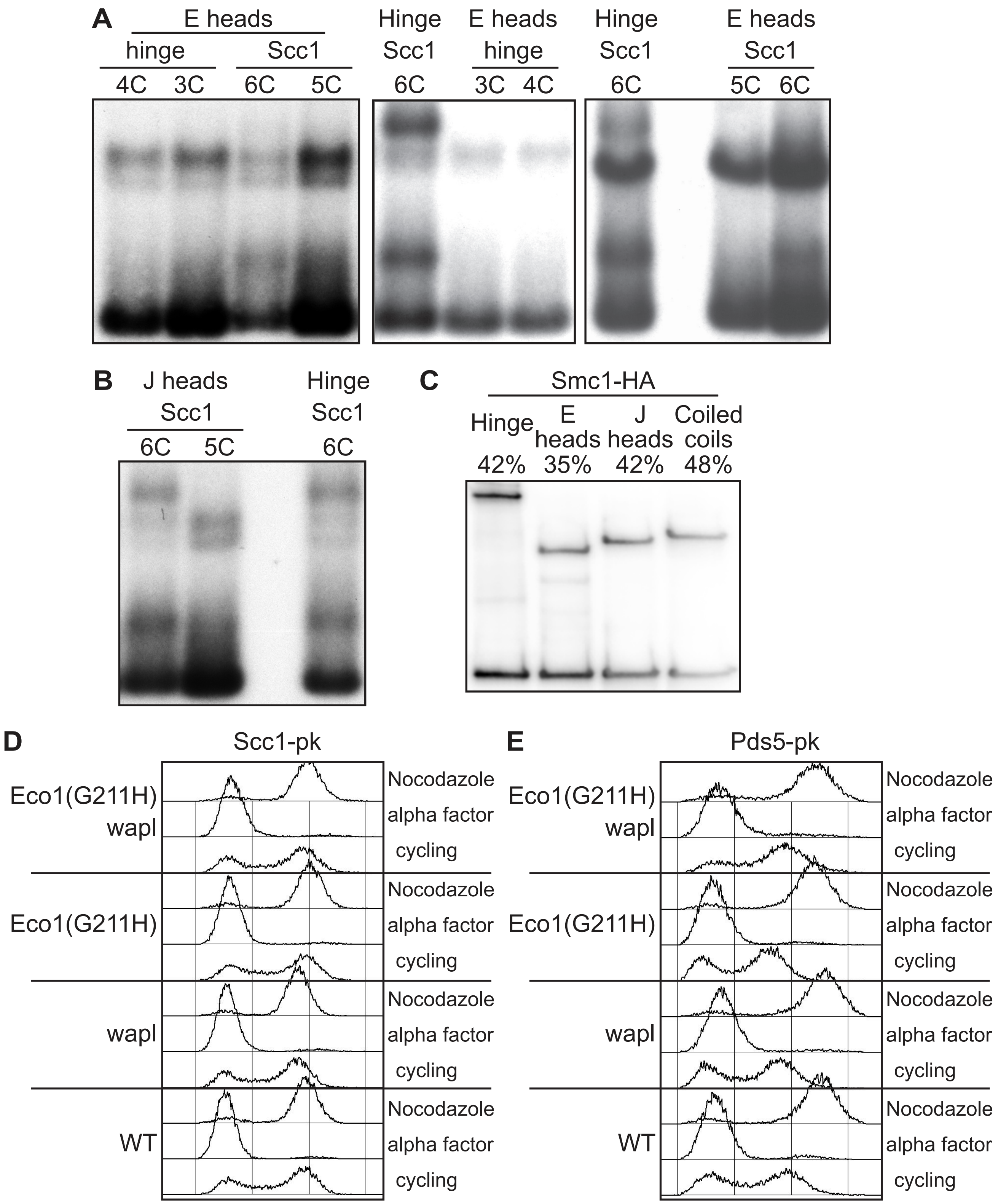

### Supplementary file 5

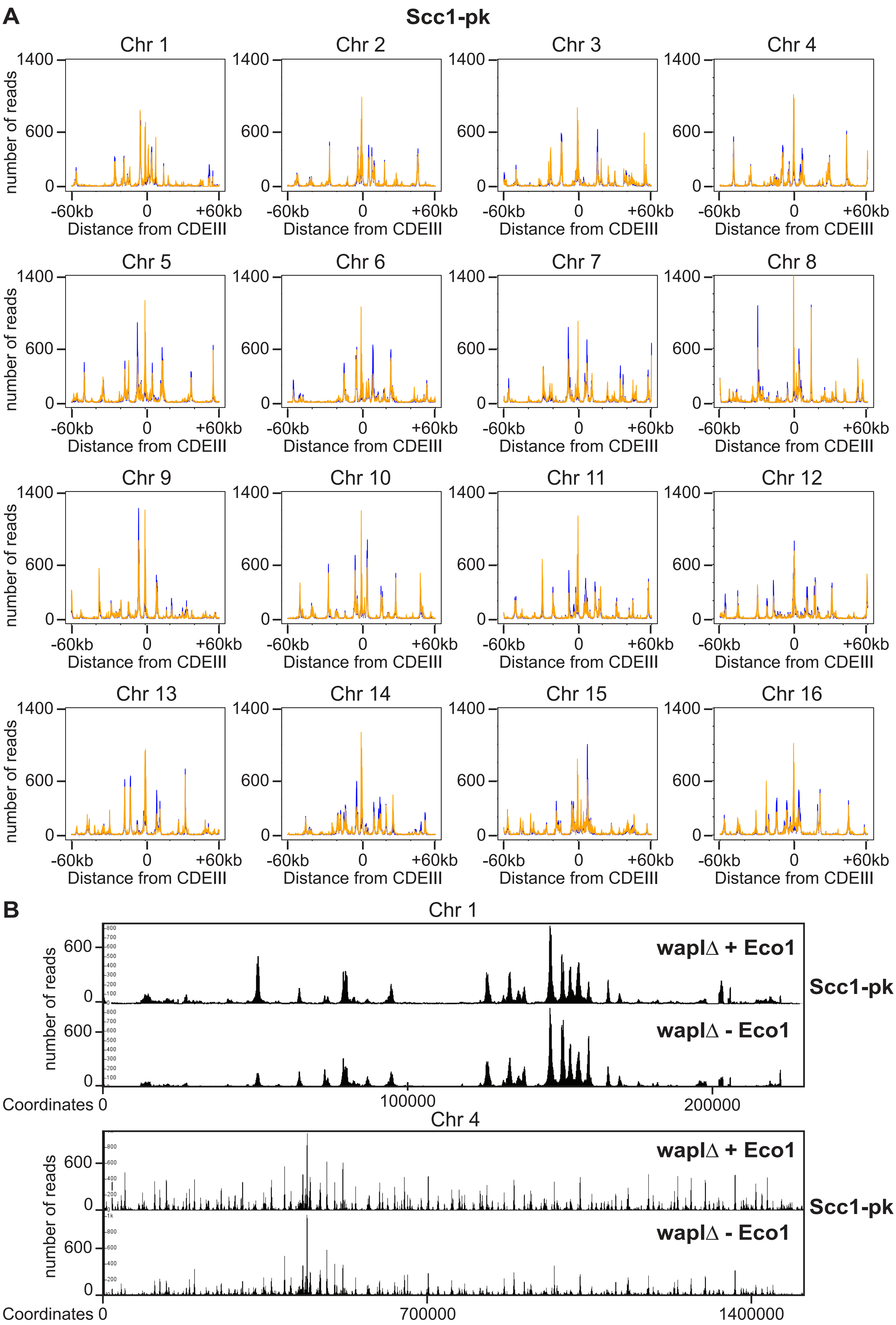

### Supplementary file 6

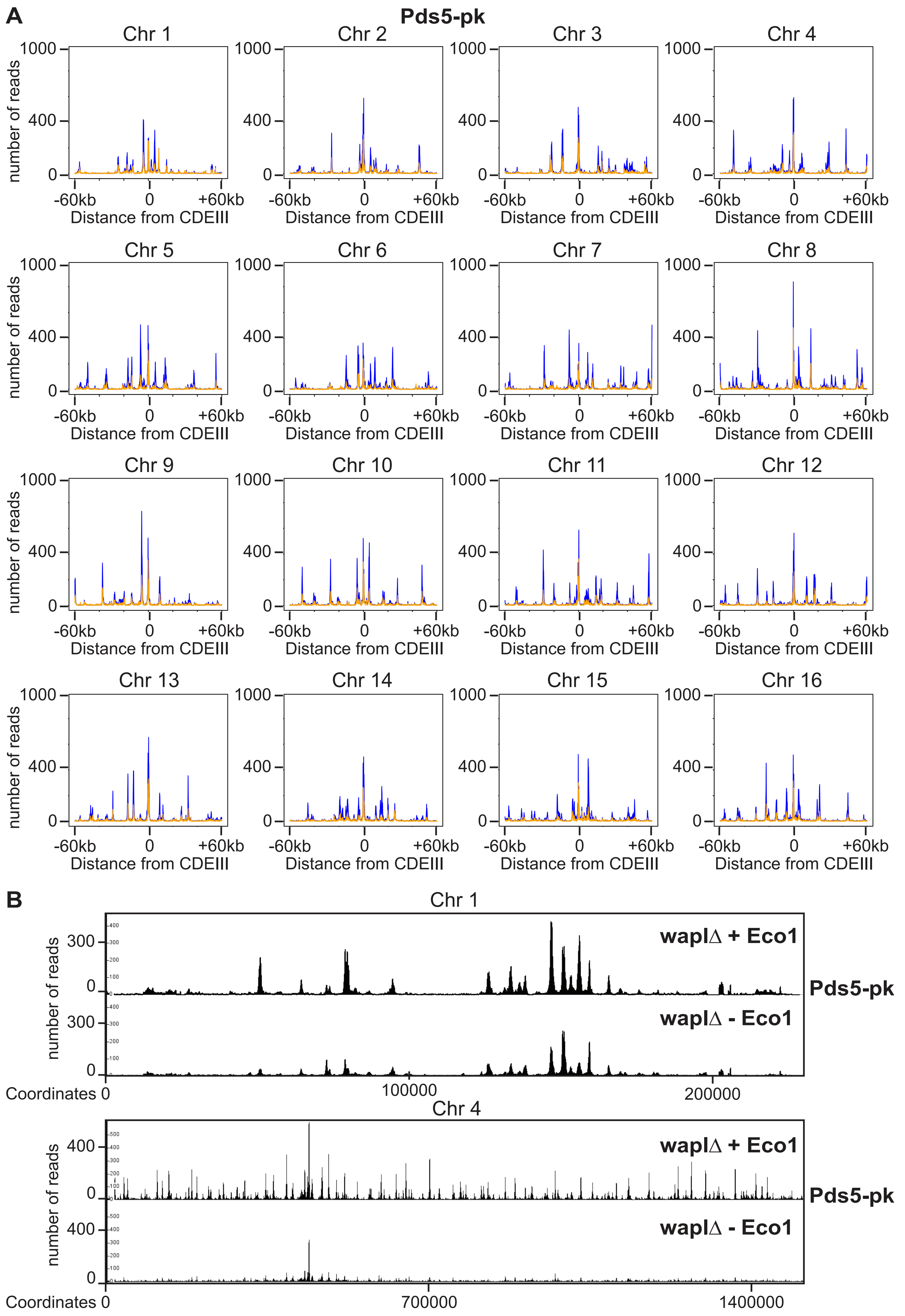
